# Evolution of embryonic *cis*-regulatory landscapes between divergent *Phallusia* and *Ciona* ascidians

**DOI:** 10.1101/371005

**Authors:** Alicia Madgwick, Marta Silvia Magri, Christelle Dantec, Damien Gailly, Ulla-Maj Fiuza, Léo Guignard, Sabrina Hettinger, Jose Luis Gomez-Skarmeta, Patrick Lemaire

**Affiliations:** Centre de Recherche en Biologie cellulaire de Montpellier (CRBM), Université de Montpellier, CNRS, Montpellier, France; Centro Andaluz de Biología del Desarrollo (CABD), Consejo Superior de Investigaciones Científicas/Universidad Pablo de Olavide/Junta de Andalucía, Sevilla, Spain; Janelia Research Campus, Howard Hughes Medical Institute, 19700 Helix drive, Ashburn, VA, USA

**Keywords:** Ascidians, embryo, chromatin, enhancer, transcription, evolution

## Abstract

Ascidian species of the *Phallusia* and *Ciona* genera are distantly related, their last common ancestor dating several hundred million years ago. Although their genome sequences have extensively diverged since this radiation, *Phallusia* and *Ciona* species share almost identical early morphogenesis and stereotyped cell lineages.

Here, we explored the evolution of transcriptional control between *P. mammillata* and *C. robusta*. We combined genome-wide mapping of open chromatin regions in both species with a comparative analysis of the regulatory sequences of a test set of 10 pairs of orthologous early regulatory genes with conserved expression patterns.

We find that ascidian chromatin accessibility landscapes obey similar rules as in other metazoa. Open-chromatin regions are short, highly conserved within each genus and cluster around regulatory genes. The dynamics of chromatin accessibility and closest-gene expression are strongly correlated during early embryogenesis. Open-chromatin regions are highly enriched in *cis*-regulatory elements: 73% of 49 open chromatin regions around our test genes behaved as either distal enhancers or proximal enhancer/promoters following electroporation in *Phallusia* eggs.

Analysis of this datasets suggests a pervasive use in ascidians of “shadow” enhancers with partially overlapping activities. Cross-species electroporations point to a deep conservation of both the *trans*-regulatory logic between these distantly-related ascidians and the *cis*-regulatory activities of individual enhancers. Finally, we found that the relative order and approximate distance to the transcription start site of open chromatin regions can be conserved between *Ciona* and *Phallusia* species despite extensive sequence divergence, a property that can be used to identify orthologous enhancers, whose regulatory activity can partially diverge.

**HIGHLIGHTS:**

- Open chromatin regions are highly enriched for *cis*-regulatory elements in ascidian early embryos.
- Ascidian Clusters of Open Regulatory Elements (COREs) are found next to regulatory genes.
- The spatio-temporal dynamics of chromatin accessibility and closest-gene expression are correlated.
- Regulatory gene expression, the *trans*-regulatory code, is conserved between distantly-related *Ciona* and *Phallusia* ascidians.
- Ascidians show pervasive use of “shadow” enhancers
- The position of homologous enhancers can be conserved, despite extensive genome divergence.

## INTRODUCTION

Development and morphogenesis are precisely orchestrated by complex Gene Regulatory Networks. These networks combine transcription factors, and the *cis*-regulatory sequences to which they bind to control gene expression. Morphological change during evolution is frequently caused by variations in the transcriptional programme, and in particular in *cis*-regulatory sequences (Stern and Orgogozo, 2008). The converse is, however, not always true. There are examples of divergent gene regulatory networks producing very similar phenotypic outputs, a scenario referred to as Developmental Systems Drift or DSD (True and Haag, 2001), exemplified by vulva specification in nematodes (Félix and Barkoulas, 2012; Sommer, 2012) or heart morphogenesis in ascidians (Stolfi et al., 2014). Our understanding of the complex relationships between genotype and phenotype currently remains too fragmentary to predict the phenotypic outcome of a regulatory mutation.

Ascidians are marine invertebrate chordates belonging to the vertebrate sister group, the tunicates (Lemaire, 2011). Like vertebrates, their embryos develop through a tadpole larval stage. Unlike vertebrates, however, ascidian embryonic development is highly stereotyped, and proceeds with invariant cell lineages. Each cell can be individually named and found across all embryos of a given species and the number, names and location of its progeny is also precisely defined (Conklin, 1905; Nishida, 1987; Guignard et al., 2018). Strikingly, even distantly related species, which diverged more than 300 million years ago share very similar or identical cell lineages. This provides a rigorous framework to compare, with cellular resolution, the evolution of developmental programmes between distantly related species. In light of this exceptional morphogenetic conservation, ascidian genomes are surprisingly divergent (Tsagkogeorga et al., 2012; Stolfi et al., 2014; Brozovic et al., 2018). Both coding and non-coding sequences evolve rapidly within the group. As a result, while non-coding sequence conservation within a genus (e. g. between *Ciona robusta* and *Ciona savignyi*, or between *Phallusia mammillata* and *Phallusia fumigata*) can be used as hallmark of functional importance, non-coding sequences across genera cannot be aligned. In addition, gene orders are frequently scrambled along chromosomes and even highly conserved gene clusters, such as the *Hox* cluster, are exploded (Duboule, 2007). Ascidians thus constitute a very interesting taxon to study DSD.

*Cis*-regulatory sequences are short (<1kb) non-coding DNA segments, which act as binding platforms for transcription factors, and control gene expression (Yáñez-Cuna et al., 2013). These sequences are so divergent between distant ascidian species that the sequences of homologous elements can generally not be aligned anymore (Oda-Ishii et al., 2005; Roure et al., 2014; Stolfi et al., 2014). In some cases, highly divergent regulatory sequences have retained the ability to respond to the same transcription factor combination (Oda-Ishii et al., 2005; Roure et al., 2014), the accumulation of *cis*-regulatory mutations thus leaving the GRN architecture intact. In other cases, however, GRN architecture have changed sufficiently for functional *cis*-regulatory sequences from one species to become “unintelligible” to another species (Lowe and Stolfi, 2018; Stolfi et al., 2014; Takahashi et al., 1999). The conservation of the function of cis-regulatory sequences between distantly related ascidians has been assayed in too few cases to estimate the relative frequency of conservation vs divergence of *cis*-regulatory activity. The relative contributions of local TF binding site turn over versus more global GRN rewiring between ascidian species also remains unknown.

In this study, we first used ATAC-seq (Buenrostro et al., 2013) to characterize the temporal patterns of chromatin accessibility in two distant phlebobranch species that diverged around 270 million years ago (Figure 1A) (Delsuc et al., 2018), *Phallusia mammillata* and *Ciona robusta*. We then used this dataset to identify in both species the early embryonic *cis*-regulatory sequences controlling the expression of ten orthologous pairs of regulatory genes. These analyses brought to light a surprising level of conservation of chromatin accessibility landscapes and *cis*-regulatory logic between these species despite extreme sequence divergence.

**Figure 1:**
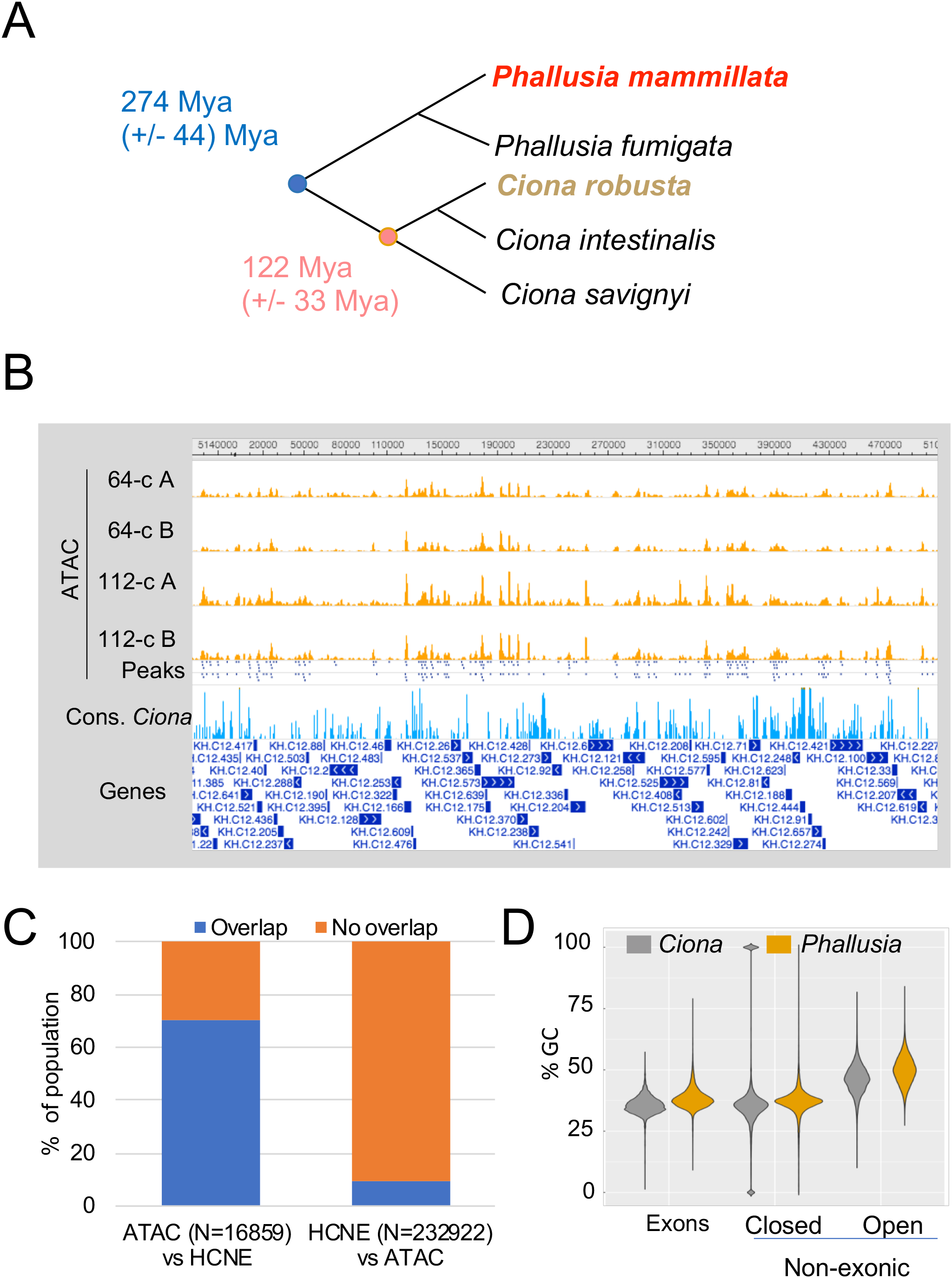
General features of chromatin accessibility in *Ciona robusta* and *Phallusia mammillata*. **A)** A cladogram with mean divergence dates and associated standard deviation (in million years from now) of the *Phallusia* and *Ciona* species discussed in this study (Delsuc et al., 2018; LN CAT-GTR + Γ4 model). **B)** Bird’s eye view of the open chromatin landscape over 500 kb of *Ciona robusta* chromosome 12 visualized with the ANISEED WashU browser. The stages at which the ATAC-seq experiments were performed and the two biological replicas (A, B) are indicated. Two rows of peaks, extracted from the union of the reads of the two replica for each stage, are shown: top, 64-cell stage; bottom, 112-cell stage. Conservation between *Ciona robusta* and *Ciona savignyi* is indicated (range 60-80 %). **C)** Overlap between chromatin accessibility (ATAC-seq peaks) and sequence conservation between *Ciona robusta* and *Ciona savignyi* within non-coding sequences. The left ATAC-seq vs HCNE bar shows the fraction of ATAC-seq peaks at the 112-cell stage overlapping a Highly-Conserved Non-coding Element: a region of LastZ alignment with more than 60% local identity. Conversely, the right HCNE vs ATAC-seq bar shows the fraction of HCNE overlapping one or more ATAC-seq peaks at the 112-cell stage. **D)** GC content analysis in *Ciona robusta* (grey) and *Phallusia mammillata* (orange) in either exons or non-exonic sequences (introns and intergenic sequences) as a function of their chromatin accessibility at the 112-cell stage (open = in an extracted ATAC-seq peak).

## MATERIAL AND METHODS

### Animal origin, *in vitro* fertilization and pharmacological treatments of *Phallusia* and *Ciona* embryos

*Phallusia mammillata* and *Ciona intestinalis* (previously *Ciona intestinalis* type B; Caputi et al., 2007) were provided by the Centre de Ressources Biologiques Marines in Roscoff, France. *Ciona intestinalis* dechorionation was done on fertilised eggs as previously described (Christiaen et al., 2009a). *Phallusia mammillata* unfertilised eggs were dechorionated in 0,1% trypsin in ASW with gentle shaking for 2 hours (Sardet et al., 2011). All live experiments were performed with dechorionated embryos. GSK3 inhibition was performed in both species by treating embryos with 4μM CHIR-99021 from the 8-cell to the stage of collection for ATAC-seq analysis.

### Gene models (names, unique gene IDs and genome assemblies)

The *Ciona robusta* genes analysed in this project were from the KH assembly (Satou et al., 2008), represented by KH2012 gene models (unique ANISEED gene IDs). For *Phallusia mammillata*, the gene models (unique ANISEED gene IDs) were from the MTP2014 assembly (Brozovic et al., 2016).

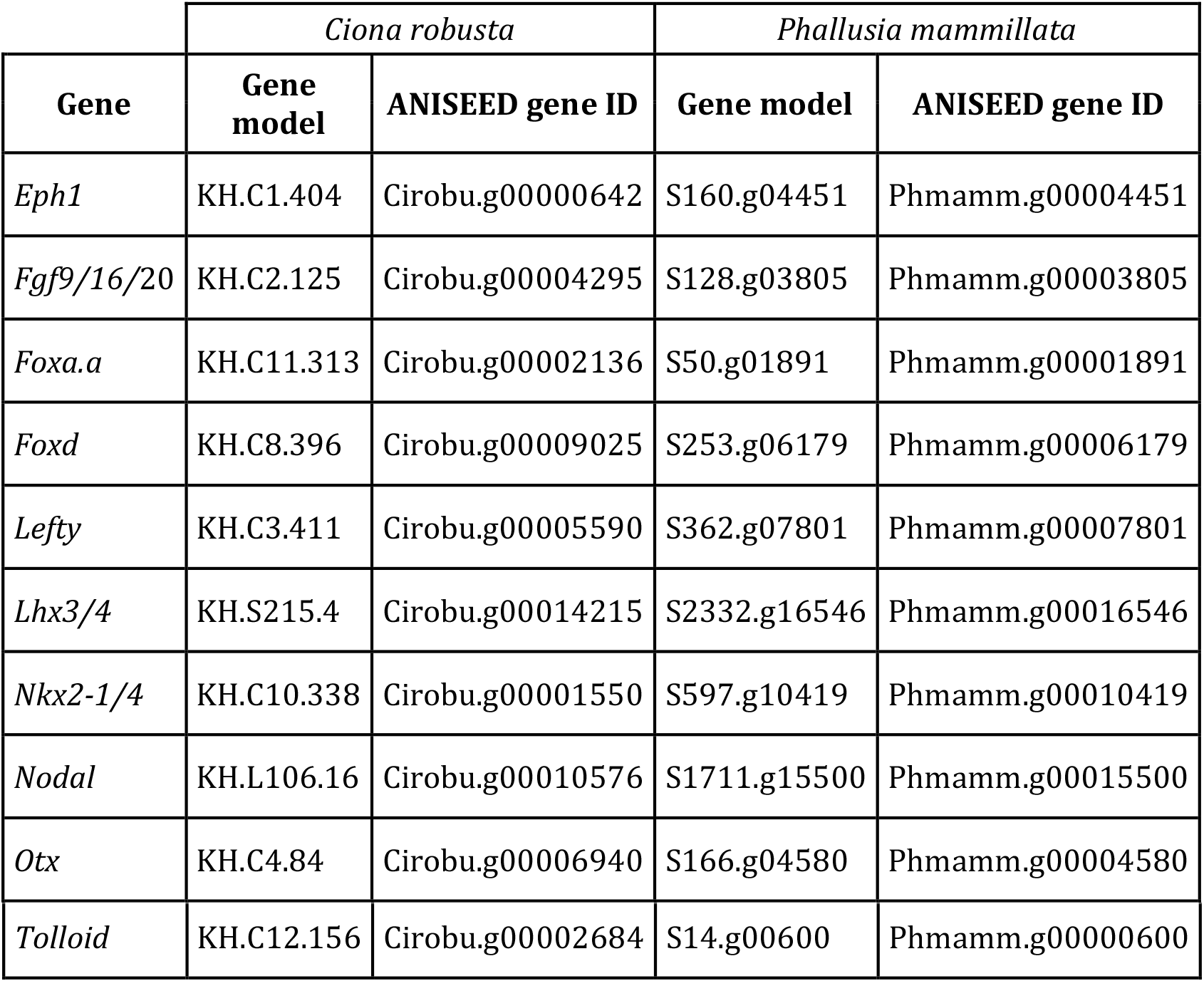

### *In situ* hybridisation

Wholemount *In situ* hybridisation was performed as previously described (Christiaen et al., 2009b). Dig-labelled probes for *Phallusia mammillata* were synthesised from the following cDNAs: *Otx* (AHC0AAA196YH07), *Lefty* (AHC0AAA31YI04), *Nkx2-1/4* (AHC0AAA267YK08), *Tolloid* (AHC0AAA14YA03), *Fgf9/16/20* (AHC0AAA22YD05), *Foxa.a* (AHC0AAA74YF05), *Foxd* (AHC0AAA228YM13), *Lhx3/4* (AHC0AAA183YA13).

The *Phallusia Nodal* and *Eph1* probes were generated by PCR from cDNA using the following primers (As in Fiuza et al., 2018): *Nodal*, Forward: 5’-CTATGGATATGACACAAGTATCGTTCTGC-3’, Reverse: 5’-GATCCTAATACGACTCACTATAGGGTTATCGACATCCACATTCT-3’; *Eph1*, Forward: 5’- C CAAC GTTGCGACTC CACTTTCACC-3’, Reverse: 5’- AC GATAAT AC GACTCACTATAGGGGTCTTGGTT AGACATCTCCCAG-3’.

### ATAC-seq assays

ATAC-seq assays were carried out essentially as described (Buenrostro et al., 2013) starting with 50-90 thousand embryonic cells, estimated from the processed volume of sedimented whole dechorionated embryos (~5nl/embryo, or 20000 embryos in 100μl). For each developmental stage, we calculated the necessary volume to reach approximately 160000 cells (16-cell stage: 50 μl, 32-cell stage: 25 μl, 64-cell stage: 12 μl, 112-cell stage: 7 μl, late gastrula: 7 μl and mid neurula: 7 μl). The non-stick 1,5ml microtubes containing the embryos were immediately placed in a microcentrifuge at 4 ºC and the embryos pelleted by spinning at 500 g for 5 min. The supernatant was carefully removed and the pellet was resuspended in 100 μl of lysis buffer, followed by about 3 minutes of vigorous pipetting. We then transferred 50 μl, 20 μl, and 10 μl of the lysate into new tubes that were centrifuged at 500 g 10 min at 4 <u>°</u>C. Meanwhile, we used the remaining 20 μl of lysate to count the nuclei using DAPI with the help of a Neubauer or Malassez chamber. This precise calculation was used to select the appropriate combination of the different aliquots (50 μl, 20 μl, 10 μl) corresponding to 50,000-90,000 nuclei. The lysis buffer was removed from all selected aliquots and the pellet(s) of nuclei were gently resuspended in 50 μl of transposition reaction mix (Nextera DNA library prep kit, Illumina, FC-121-1030). If more than one aliquot were needed to reach the appropriate cell number, they were pooled in a final volume of 50 μl of mix. The transposition reaction was incubated for 30 min at 37 °C. Immediately after transposition, we purified DNA with the Qiagen MinElute Kit (Qiagen, 28004) following the manufacturer’s instructions. Purified DNA can be stored at - 20°C or immediately processed to generate the ATAC-seq library. To generate the ATAC-seq library, each sample was subjected to 15 cycles of PCR amplification using the primers and conditions described in Supp. Table 1 of (Buenrostro et al., 2013). We then purified the amplified library using Qiagen MinElute Kit (Qiagen, 28004) and determined its concentration with the Qubit dsDNA BR Assay Kit (Molecular Probes #Q32850) according to the manufacturer’s instructions. A minimum concentration of 30 ng/mL is preferred for NGS. The quality of the amplified library was determined by running an aliquot on a 2% agarose gel. A smear covering 200 bp to 1 kb should be observed. Depending on the quality of the samples, bands corresponding to mono-(200 bp) and di-nucleosomes (400 bp) were observed. Libraries were stored at -20°C. Sequencing was done using an Illumina HiSeq4000 sequencer (BGI) and approximately 20 million paired 100b sequence reads were obtained per sample.

### ATAC-seq sequence pre-processing, alignment and peak calling

Illumina adapters were trimmed with Cutadapt (Martin, 2011). Alignment on the *Phallusia mammillata* and *Ciona robusta* genomes was performed with bowtie2, (very-sensitive parameter) (Langmead and Salzberg, 2012). Although our *Ciona* dataset came for *Ciona intestinalis* animals, the very poor quality of the *Ciona intestinalis* genome assembly (Brozovic et al., 2016) led us to align these sequences onto the closely related *Ciona robusta* genome. In both *Phallusia* and *Ciona*, only a minority of sequences aligned to the nuclear genome (*Ciona* ~30%; *Phallusia* ~17%). Significant Peaks were extracted with MACS2 (Zhang et al., 2008) with the following parameters for *Phallusia mammillata*: gsize=187138625 (genome size = genome mappable size excluding repeated sequences) –nomodel -f BAMPE -q 0.01; for *Ciona*: gsize= 99000000 –nomodel -f BAMPE -q 0.01. *Ciona* and *Phallusia* raw ATAC-seq sequences were submitted to the Short Read Archive (Bioproject IDs <u>PRJNA474983</u> and <u>PRJNA474750</u>, respectively). Peaks and short read coverage normalized per million of aligned reads can be seen on the *<u>Ciona robusta</u>* and *<u>Phallusia mammillata</u>* ANISEED WashU genome browser (in the “whole-organism epigenomics data” public track Hubs).

### Construction of a reference peak dataset for each species

To compare chromatin accessibility between experimental conditions and developmental stages, we constructed a reference set of non-coding non-overlapping ATAC-seq peaks for each species. We started from MACS-called peaks for each biological sample and excluded peaks overlapping exonic sequences. A chromatin accessibility score for each reference peak was defined by allocating for each condition the sum of the depth of normalized ATAC-seq sequence coverage over all peak bases (all accessibility score values below 1 were considered unreliable and arbitrarily set to 1 to calculate fold changes). This score was used to select for each condition the peaks found in the top 2/3 of this accessibility score distribution (*Phallusia*: ATAC score >500; *Ciona*: ATAC score >300), thereby excluding smaller peaks more influenced by experimental noise.

The reference peak set for each species was built by merging overlapping peaks called in individual conditions, only retaining peaks found on a scaffold harbouring one or more coding genes. This generated reference sets of 15170 and 15029 peaks in *Ciona* and *Phallusia*, respectively.

We then associated to each peak the gene, whose 5’-most transcript model was closest to the centre of the ATAC-seq peak, a procedure which manual examination of more than 100 peaks revealed as accurate in more than 80% of cases, the majority of errors coming from unsatisfactory gene models (truncated or missed genes). The *Ciona* and *Phallusia* reference peak sets were associated to 6644 and 7577 coding genes respectively, and we recorded the position of the peak with respect to the gene body. Peaks were sorted into 5 categories: intronic and intergenic <1kb, 1-2kb, 2-5kb, >5kb from the start of the most 5’- extending transcript model. Expression data, normalized as fpkm were downloaded from ANISEED. All fpkm values below 1 were considered unreliable and arbitrarily set to 1 for the calculation of fold-changes.

### Gateway cloning and electroporation of the regulatory sequences

Constructs to test the activity of candidate regulatory sequences from both species were generated by Gateway cloning into the pSP1.72.RfB::NLS-LacZ destination vector (Lamy et al., 2006), or into a modified version, in which the *Foxa.a* minimal promoter from *Ciona robusta* (ANISEED ID: REG00001035) was placed upstream of NLS-LacZ to generate pSP1.72.RfB.pFoxa.a::NLS-LacZ, or into pSP1.72.RfB.pFog::NLS-LacZ as indicated in Table S3. The primers for each of the tested sequences, the ANISEED IDs of their constructs and their spatio-temporal patterns of transcriptional activity and reference in the ANISEED database are listed in Tables S3 and S4. The genomic DNA used to amplify by PCR the genomic fragments from *Ciona robusta* was a kind gift from Dr. A. Spagnuolo (Naples). *Phallusia mammillata* genomic DNA was extracted from animals collected in the Roscoff area.

Electroporations were performed as previously described (Bertrand et al., 2003), with a single pulse of 16ms at 50V for *Ciona intestinalis* and two pulses of 10ms at 40V for *Phallusia mammillata*. X-gal staining was performed as previously described (Bertrand et al., 2003). Sequences that drove activity of the LacZ reporter gene in fewer than 10% of electroporated embryos or larvae, or only in the mesenchyme (known to be a tissue displaying ectopic staining for many sequences) were considered inactive. More than 100 embryos were counted for each construct and experiment.

## RESULTS AND DISCUSSION

### Regions of open chromatin in ascidian early embryos are enriched in *cis*-regulatory sequences

To characterize the *cis*-regulatory landscapes during early ascidian embryogenesis, we first mapped open chromatin regions using ATAC-seq (Buenrostro et al., 2013) on whole *Ciona* and *Phallusia* embryos at the 64- and 112-cell stages and identified regions of open chromatin using the MACS2 peak caller (Zhang et al., 2008). Comparison of the biological replicas indicated that the process was highly reproducible (Figure 1B).

In *Ciona*, MACS2 identified 24574 (median length 119 bp) and 19397 (median length 189 bp) non-coding ATAC-seq peaks at the 64- and 112-cell stages, respectively, covering 3.32% and 4.38% of the genome sequences. As expected, most (70%) open chromatin regions were located in regions of high (>60%) sequence identity between *C. robusta* and *C. savignyi* estimated to have diverged around 120 million years ago (Delsuc et al., 2018) (Figure 1A, C). Conversely, only around 10% of non-coding sequences with more than 60% of identity between *C. robusta* and *C. savignyi* overlapped with open chromatin regions during early development (Figure 1C). The majority of non-coding conserved sequences may thus correspond to *cis*-regulatory sequences acting much later during development or in adults. In *Phallusia*, similarly, MACS2 identified 10632 (median length 219 bp) and 14240 (median length 123 bp) non-coding ATAC-seq peaks, covering 1.27% and 0.93% of the assembled genome (Brozovic et al., 2016) at the 64- and 112cell stages, respectively. In both *C. robusta* and *P. mammillata*, the GC content of ATAC-seq peaks was higher than that of other non-exonic and exonic sequences (Figure 1D), consistent with previous findings on *cis*-regulatory sequences in other model organisms (Li et al., 2007).

Thus, as in other animal taxa, open chromatin regions detected by ATAC-seq during early ascidian embryogenesis cover a small part of the non-coding genome, enriched in GC and highly-conserved within the *Ciona* genus.

### Ascidian regulatory genes are surrounded by Clusters of Open *cis*-regulatory Elements (COREs)

In vertebrates, open regulatory elements are unevenly distributed along the chromosomes. They show a higher density around developmental regulators, defining Clusters of Open *cis*-regulatory Elements (COREs) (Gaulton et al., 2010), which have been associated to the super-enhancers controlling the expression of genes controlling cell fate decisions (Pott and Lieb, 2015).

We found likewise that at the 64- and 112-cell stages, non-coding ATAC-seq peaks were unevenly distributed along the genomes of both ascidian species, with clustered peaks separated by regions of much lower density (Figure 2A). 33% and 65% of *Phallusia* and *Ciona* ATAC-seq peaks defined at these stages belonged to 821 and 2099 clusters of at least 5 ATAC-seq peaks within 10kb, respectively. Genes overlapping with or immediately adjacent to such clusters were highly enriched for molecular functions linked to transcription and signalling and for biological processes linked to development, regulation, transcription and signalling (Figure 2B).

**Figure 2:**
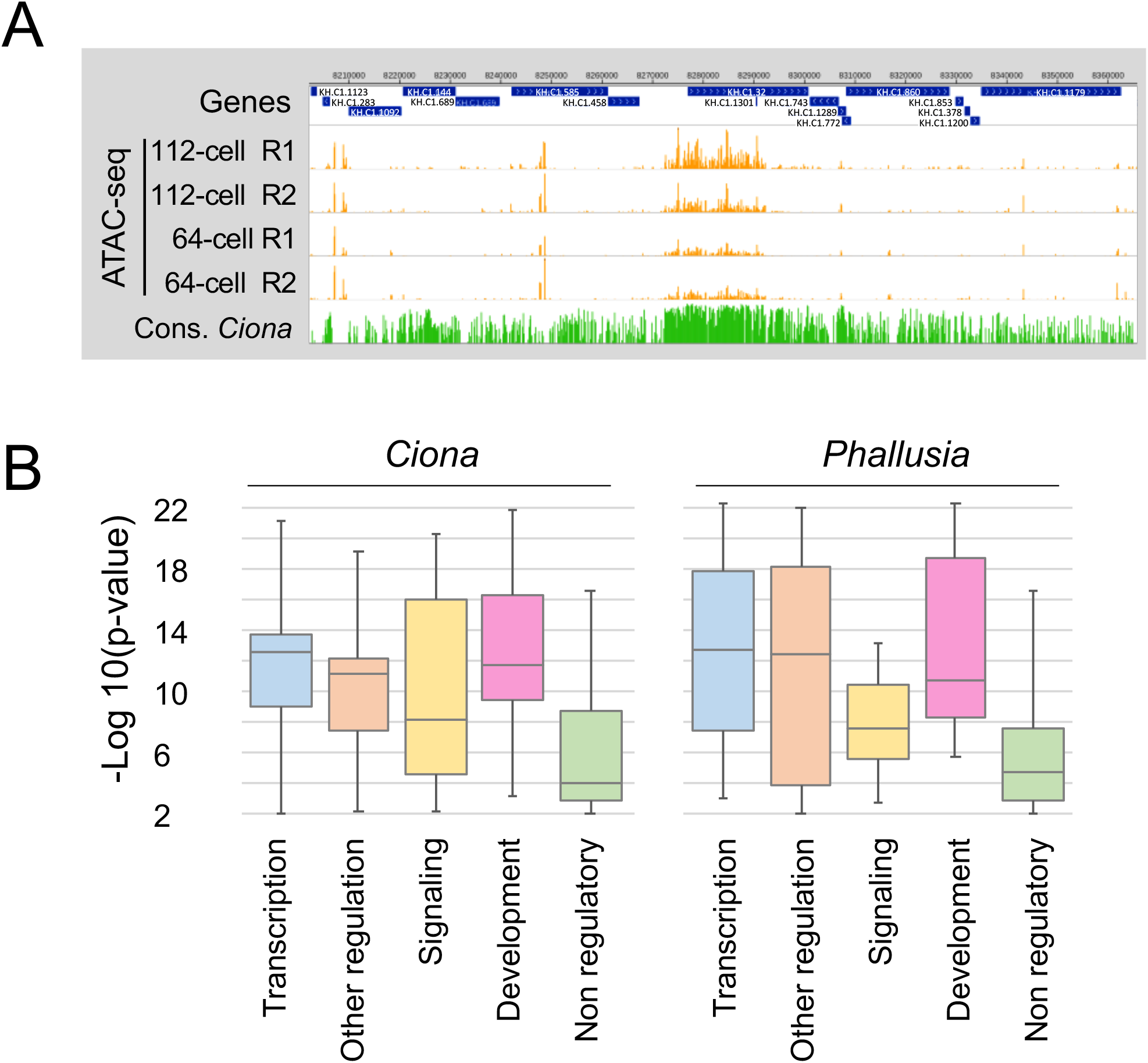
Clusters of Open Regulatory Elements (COREs) are found in the vicinity of ascidian developmental regulatory genes. **A)** A bird’s eye view of 150kb of *Ciona robusta* chromosome 1 showing the clustering of ATAC-seq signal around gene KH.C1.32, a C2H2 zinc finger gene. The ATAC-seq signal at the 64-cell and 112-cell stages and the conservation between *Ciona robusta* and *Ciona savignyi* (identity >60%) are shown. **B)** Most-enriched GO terms -(Log10(P-value)>2, Terms describing Molecular Function or Biological Processes only) in genes overlapping or immediate neighbours to COREs (≥5 noncoding peaks within 10kbs) in both species. Each enriched term was manually assigned to one of the 5 shown categories. Note the poorer enrichment of “Non-regulatory” terms (P<0.005 with any other class, t-test). GO term enrichments were computed from the Bingo tool of Cytoscape (Maere et al., 2005) from ANISEED GO annotations.

Thus, ascidian embryos, despite their simpler anatomy, reduced cell numbers and compact genomes share with vertebrates the presence of conserved clusters of open chromatin regulatory elements (COREs) around developmental regulatory genes.

### Construction of a reference set of open chromatin regions during early ascidian embryogenesis

To assess the spatio-temporal dynamics of chromatin accessibility in *Phallusia* and *Ciona*, we extended our *Phallusia* dataset to encompass the 16-cell, 32-cell, 64-cell, 112-cell, mid-gastrula and mid-neurula stages. The *Ciona* dataset was also extended with mid-gastrula and mid-neurula stage samples. In addition, to estimate the impact of tissue specification on chromatin accessibility landscapes, we collected in both species ATAC-seq data from early gastrula embryos in which the GSK3 kinase had been pharmacologically inhibited from the 8-cell stage, a treatment known to activate the canonical Wnt/ß-catenin signalling pathway, thereby converting most embryonic cells into endoderm (Imai et al., 2000).

To compare the chromatin accessibility landscapes more quantitatively across conditions and species, and to relate them to patterns of gene expression, we compiled a reference set of around 15000 noncoding, non-overlapping peaks for each species including all peak regions called across all conditions (see methods). Only peaks with a chromatin accessibility score in the top 2/3 of the score distribution in at least one condition were kept. We then associated to each peak its closest coding gene, its position with respect to this gene and the expression pattern of this gene at the 64-cell, early gastrula, mid-neurula and mid tailbud stages (see Tables S1 and Table S2 for *Ciona* and *Phallusia*, respectively). All ATAC-seq datasets generated in the frame of this project can be explored through public hubs in the ANISEED *Ciona robusta* and *Phallusia mammillata* WashU. Raw reads were deposited in the NCBI Short Read Archive (Bioprojects <u>PRTNA474750</u> and <u>PRTNA474983</u>).

### Temporal dynamics of chromatin accessibility in *Phallusia mammillata* and its relationship to gene expression

Starting from this dataset, we first analysed the temporal dynamics of chromatin accessibility between the 64-cell and mid-neurula stages and its relationship to gene expression. Of the reference peaks whose accessibility scores were in the top 2/3 at the mid-neurula or 64-cell stages, approximately 50% showed less than a 2-fold variation in their accessibility between the 64-cell and mid-neurula stages (Figure 3A). These results are in keeping with *Drosophila, C. elegans* and vertebrates in which a minority of loci show a temporally dynamic chromatin accessibility in embryos (Cusanovich et al., 2018; Daugherty et al., 2017; Hagey et al., 2016; Liu et al., 2018; Preissl et al., 2018; Thomas et al., 2011; Wu et al., 2016). We note however that approximately 50% of the peaks showed at least a 2-fold temporal variation in the pattern of chromatin accessibility, as exemplified for the Matrilin-related gene (Figure 3B).

**Figure 3:**
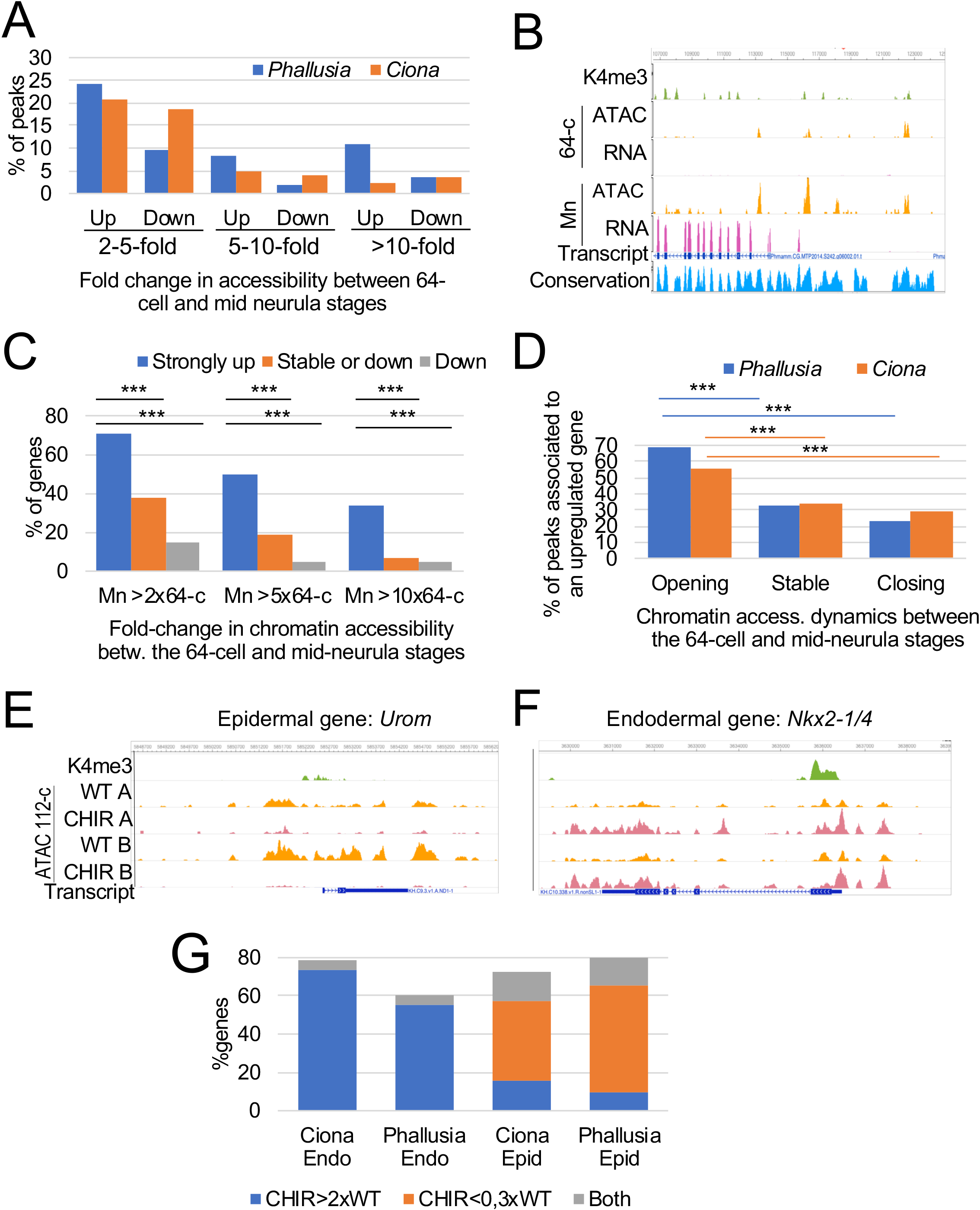
Relationship between spatio-temporal patterns of chromatin accessibility and gene expression in *Phallusia* and *Ciona*. **A)** Fraction of Phallusia reference ATAC-seq peaks with the indicated temporal dynamics between the 64-cell and mid-neurula stages. “Up” (“Down”) means that chromatin accessibility increased (decreased) between the 64-cell and mid-neurula stages, respectively. **B)** View of the chromatin accessibility (normalized ATAC-seq signal) and RNA expression (normalized RNA-seq signal) around the *Matrilin-like* gene (Phmamm.CG.MTP2014.S242.g06002). ChIP-seq results H3 K4me3 mark are from (Brozovic et al., 2018). The Conservation track shows the local sequence conservation between *Phallusia mammillata* and *fumigata*. ATAC, RNA: ATAC-seq and RNA-seq signals at the indicated stage (64-cell, mid-neurula). **C)** Percentage of *Phallusia mammillata* genes associated to at least one ATAC-seq peak with increasing chromatin accessibility between the 64-c and mid neurula stage as a function of their expression dynamics. Strongly up: Max fpkm (Neurula/64-c, Tailbud/64-c, Neurula/e. Gastrula, Tailbud/ e. Gastrula)>10 (N= 384). Stable or down: Max fpkm over same stages ≤1 (N=92). Down: Max fpkm over same stages <0,3 (N=40). The analysis is restricted to genes with low (fpkm<2) maternal expression. Mn>2×64-c indicates that the accessibility score is twice as high at the mid-neurula stage than at the 64-cell stage. ***: P≤0.0002 “N-1” Chi-squared test. **D)** Percentage of peaks associated to a strongly (>10-fold) upregulated gene in *Ciona* and *Phallusia*, as a function of the temporal dynamics of chromatin accessibility. Fold change of ATAC-seq score between the 64-cell and mid-neurula stages: Opening: >5-fold change (Phallusia N= 447, *Ciona* N= 229); Stable: 0,9-1,1-fold change (*Phallusia* N= 111, *Ciona* N= 114); Closing: <0,5 fold change (*Phallusia* N= 89, *Ciona* N= 98). ***: as in C. **E, F)** View of the chromatin landscapes around the *Ciona* Nkx2-1(KH.C10.338) and Urom (KH.C9.3) genes. The normalized ATAC-seq signals for the two replicas A and B at the 112-cell stage in Wild-Type (WT) and GSK3-inhibited embryos (CHIR) are shown. **G)** Changes in chromatin accessibility around predominantly endodermal (N=23) and epidermal (N=26) *Ciona* genes and their *Phallusia* 1-to-1orthologs (N=20 for each category) (genes listed in the “Comments” column of Tables S1 and S2) in Wild-Type (WT) and GSK3-inhibited (CHIR) 112-cell embryos in *Phallusia* and *Ciona*. CHIR>3xWT: association to peaks of 2fold increased accessibility upon CHIR treatment; CHIR<0,3xWT: association to peaks of 3-fold decreased accessibility upon CHIR treatment; Both: association to peaks of increased and decreased accessibility.

To address a possible correlation between the temporal dynamics of gene expression and chromatin accessibility in both species, we used published time-courses of RNA-seq expression levels in both species (Brozovic et al., 2018). We first selected three sets of genes based on their temporal dynamic of expression: a set of genes upregulated at least 10-fold between the 64-cell and mid-tailbud stages; a set of genes not upregulated; and a set of at least 3-fold downregulated genes over this time window. We then computed the temporal evolution of chromatin accessibility of the peaks associated to these genes. As shown for *Phallusia* on Figure 3C, 70%, 50% and 33% of upregulated genes between the 64-cell and mid-tailbud stages were associated to peaks with more than a 2-, 5- and 10-fold increased accessibility at the neurula stage. By contrast, only 15%, 5% and 5% of downregulated genes were associated to such peaks.

Conversely, we selected peaks with 5-fold increased accessibility, stable accessibility and 2-fold decreased accessibility between the 64-cell and mid-neurula stages and analysed the temporal dynamics of gene expression of the associated genes. We restricted this analysis to peaks associated to genes with low maternal expression (fpkm<2), as maternal expression can mask the temporal dynamic of zygotic gene expression. In both species, a majority of peaks showing increased accessibility at the mid-neurula stage were associated to genes strongly upregulated between the cleavage and tailbud stages (Figure 3D). This was not the case for peaks showing decreased accessibility over time.

We conclude that although only a minority of ATAC-seq peaks show temporal regulation during early embryogenesis, a majority of temporally upregulated genes are associated to peaks with increased local accessibility, while a majority of peaks with increased accessibility over time are associated to upregulated genes. In many cases, the temporal dynamics of chromatin accessibility in the vicinity of a gene can thus be used as a predictor of its temporal expression dynamics.

### Relationship between fate specification and chromatin accessibility

To estimate the impact of tissue specification on chromatin accessibility landscapes, we compared the ATAC-seq signals at the early gastrula stage between wild-type embryos and embryos enriched in endodermal precursors following treatment with the CHIR-99021 GSK3 inhibitor from the 8-cell stage. GSK3 inhibition led to a drastic reduction of chromatin opening in the vicinity of the *Urom* gene, expressed in early gastrula epidermal precursors (Figure 3E) and a converse increased accessibility around the *Nkx2-1/4* gene, expressed in endodermal precursors (Figure 3F).

*The Ciona robusta* (formerly *Ciona intestinalis* type A; Caputi et al., 2007) expression profiles of many genes, including most transcription factors and signalling molecules have previously been characterised by *in situ* hybridisation (Imai et al., 2006, 2004). To more generally assess the correlation between tissue-specific gene expression and chromatin accessibility, we mined the ANISEED *in situ* hybridization data for *Ciona* genes expressed mostly in either epidermal or endodermal precursors during early embryogenesis. This search returned 26 and 23 genes, respectively. Figure 3G illustrates that endodermal genes were predominantly associated to ATAC-seq peaks with increased chromatin accessibility upon CHIR treatment, while conversely epidermal genes were primarily associated to ATAC-seq peaks of decreased accessibility upon GSK3 inhibition. This analysis suggests that fate specification and the dynamics of chromatin accessibility also seem to be strongly correlated during ascidian early embryogenesis.

### Conservation of early regulatory gene expression between *Ciona robusta* and *Phallusia mammillata* up to the early gastrula stage

While extensive expression datasets are available for *Ciona robusta*, much fewer data are available for *Phallusia mammillata*. Using the ANISEED orthology section, we identified the *Phallusia mammillata* orthologues of the 49 *Ciona robusta* epidermal and endodermal genes studied in the previous section. The *Ciona* endodermal and epidermal gene sets had each 20 1-to-1 orthologs in *Phallusia mammillata*. Interestingly, chromatin accessibility landscapes around *Phallusia* orthologs genes matched the behaviour seen for the *Ciona* orthologs, suggesting a conservation of chromatin accessibility landscapes despite extensive sequence divergence.

The similarity of the chromatin accessibility profiles in WT and CHIR-treated embryos suggests a conservation of the expression pattern of *Phallusia* orthologues of *Ciona* genes. We thus set out to determine by *in situ* hybridization the spatial and temporal expression patterns up to the early gastrula stage of the *Phallusia* orthologs of the 1-to-1 orthologues of four known regulators of *Ciona* endoderm development, *Foxa-a, Fgf9/16/20, Lhx3/4* and *Nkx2-1/4* (Hudson et al., 2016; Ristoratore et al., 1999; Satou et al., 2001) and 5 regulatory genes expressed in endoderm precursors but whose function in endoderm development remains hypothetical, *Nodal, Otx, Tolloid, Lefty*, and *Eph1*. This set was completed with the unique *Phallusia* orthologue of a recently-duplicated *Ciona* endoderm specifier, *Foxd*, as the two *Ciona* paralogs are so similar (96% identity at the nucleotide level) that the published *in situ* hybridisation profiles may reflect the combined expression of both genes.

Figure 4A presents the expression patterns of these 10 genes in *Phallusia* and *Ciona*. Expression patterns were scored in a qualitative manner: based on the staining pattern, each cell was considered to either express or not a given gene. Thanks to the conservation of early cell lineages between *Ciona* and *Phallusia*, the Boolean expression patterns of orthologs could be compared with a cellular level of resolution. This set of *in situ* hybridisation experiments may overestimate divergence as their interpretation depends on the duration of the alkaline phosphatase enzymatic staining (Figure 4B). For instance, at the 64-cell stage, a short staining time with an *Fgf9/16/20* probe in *Phallusia mammillata* detects expression in the cells where the *Ciona Fgf9/16/20* ortholog was previously described (A7.4, A7.8 and B7.4) (Imai et al., 2004), while a longer staining detected additional weak *Phallusia Fgf9/16/20* expression in A7.1, A7.2, A7.5 and A7.6, in which no expression was reported in Ciona. A similar situation was also found in the case of *Lefty* at the 64-cell stage.

**Figure 4:**
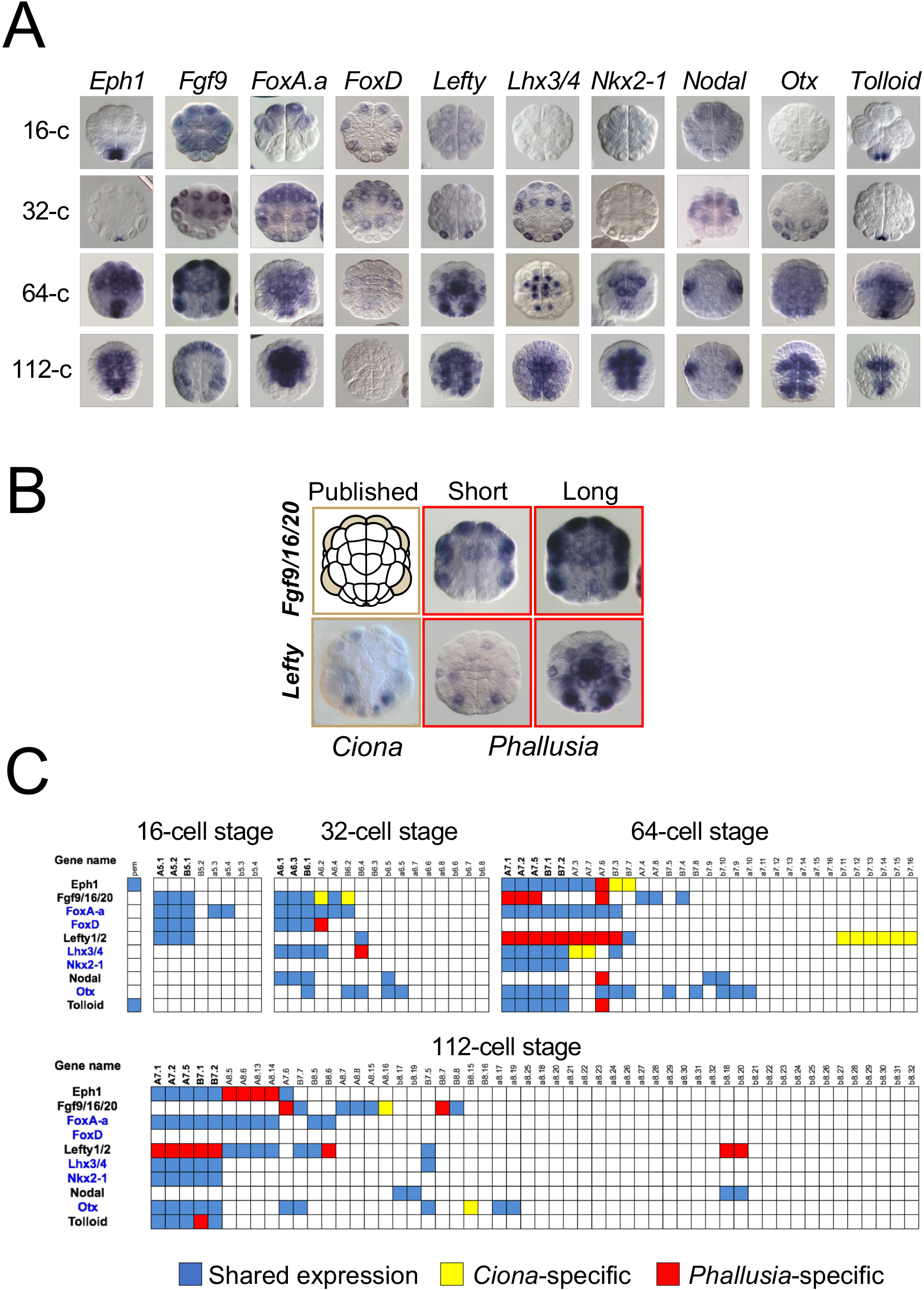
Conserved temporal and spatial early expression profiles between *Ciona* and *Phallusia* regulatory genes. **A)** *In situ* hybridisation in *Phallusia mammillata* of the indicated genes at the 16-, 32, 64- and 112-cell stages. Anterior is to the top. All of the images show vegetal views except for *Foxa.a* and *Lefty* at the 16-cell stage, which show an animal view. *Fgf9/16/20* is abbreviated as *FGF9*. **B)** Example *in situ* hybridizations for two genes, *Lefty* and *Fgf9/16/20* at the 64-cell stage in the published literature (*Ciona robusta*, Imai et al., 2004) and in our *Phallusia mammillata* experiments, showing the impact of the short or long duration of the NBT/BCIP staining reaction on the interpretation of expression patterns. **C)** Comparison table of ortholog gene expression at the cellular level at the 16-, 32-, 64- and 112-cell stages. Transcription factor genes are indicated in blue. Cell identities are indicated on top of each column. Endoderm precursor identities are in bold. Blue cells: shared expression between *Ciona robusta* and *Phallusia mammillata*; red cells: *Phallusia*-specific expression; yellow cells: *Ciona*-specific expression.

Despite this potential caveat, conservation of gene expression between the two species was overall high ((Figure 4C). It varied from perfect cell-to-cell conservation of the expression pattern (*Nkx2-1/4* and *Foxa.a*), minor divergence (expression status differs in up to 15% of expressing cells: *Otx, Tolloid, Nodal, Lhx3/4, Foxd*), and more significant divergence (expression status differs in 35 to 65% of expressing cells: *Eph1, FGF9/16/20, Lefty*).

We conclude that the early expression patterns of a majority of our ten test genes are well conserved between *Ciona* and *Phallusia*, some of the observed differences possibly reflecting limitations of the colorimetric *in situ* hybridisation protocol we used. Future work based on single-cell RNA seq in ascidians (Horie et al., 2018; Sharma et al., 2018; Treen et al., 2018; Wang et al., 2017) may be required to discriminate technical issues from *bona fide* evolutionary changes. In the following sections, we used our extensive ATAC-seq dataset to identify the *cis*-regulatory sequences driving the conserved expression of these genes and to compare their activity in both species.

### Most open chromatin regions have embryonic *cis*-regulatory activity

To confirm the helpfulness of our ATAC-seq dataset to identify the *cis*-regulatory sequences driving the expression of the ten ortholog pairs in both species, we first surveyed the chromatin state of known *Ciona* elements controlling the expression of four of these genes. Figure 5A illustrates that open chromatin regions around the *Ciona Otx* gene overlapped with previously characterized *cis*-regulatory regions and with ChIP-seq peaks for *GATAa*, a major regulator of *Otx* at the 32-cell stage. Five previously identified *Ciona robusta cis*-regulatory sequences controlling the *Fgf9/16/20* (ANISEED REG00001080, Figure 4B, Oda-Ishii et al., 2016), *Foxd* (ANISEED REG00001115, Oda-Ishii et al., 2016), *Lefty* (ANISEED REG00001118, Oda-Ishii et al., 2016) and *Nkx2-1/4* (ANISEED REG00000321, Fanelli et al., 2003) genes in early mesendodermal territories also mapped to open chromatin regions (not shown).

**Figure 5:**
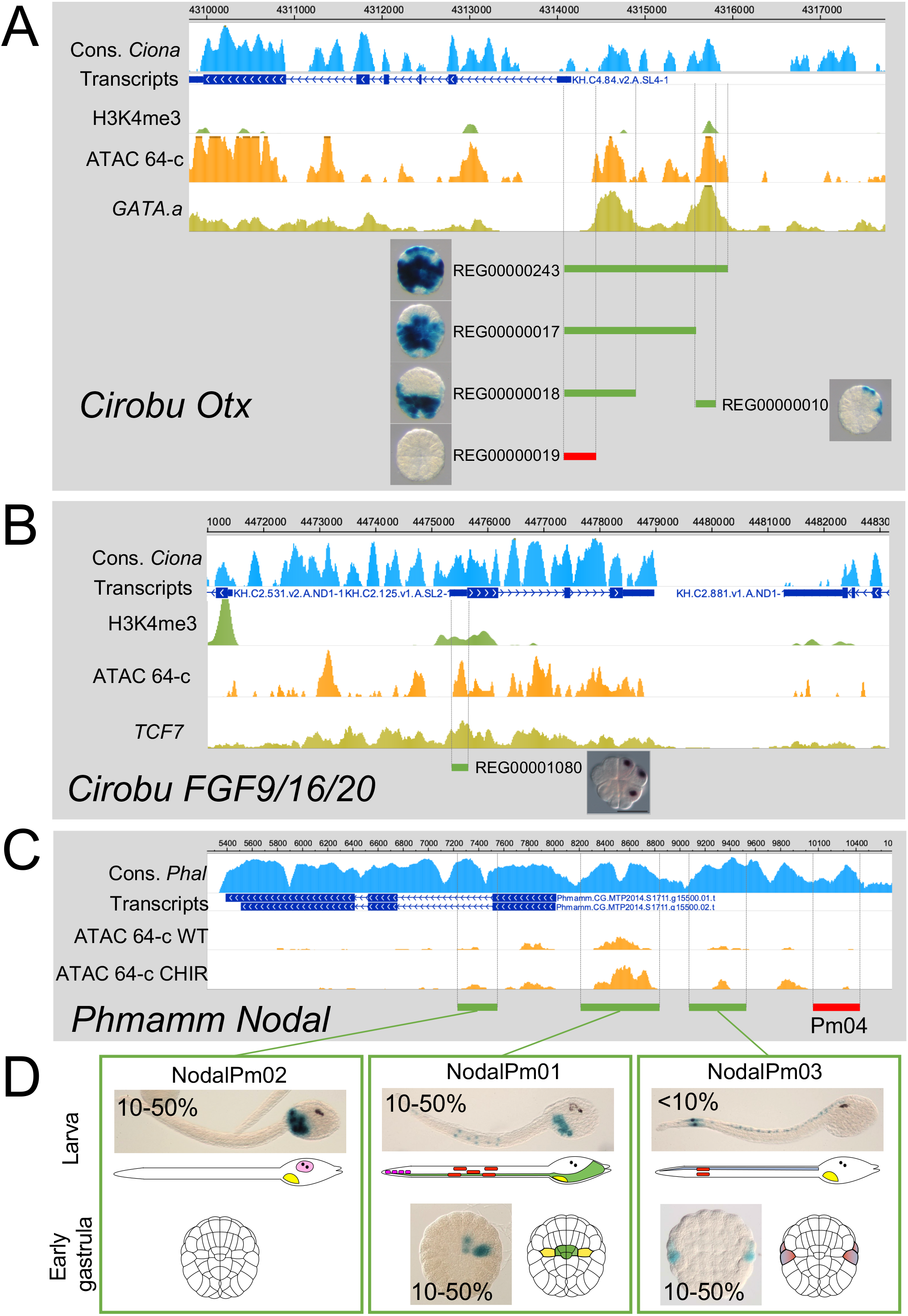
Relationship between chromatin accessibility and *cis*-regulatory activity in *Ciona* and *Phallusia*. **A)** Position of previously published *Ciona robusta* regulatory sequences for *Otx* (Bertrand et al., 2003) with respect to ATAC-seq signal (orange) at the 64-cell stage, of Gata.a ChIP-seq signal (yellow) at the 32-cell stage (Oda-Ishii et al., 2016) and of H3K4me3 ChIP-seq signal (green) at the early gastrula stage (Brozovic et al., 2018). Published pictures of representative embryos are shown next to the tested constructs. The profile of *Ciona robusta/Ciona savignyi* sequence identity (>60%) is indicated in blue. The ANISEED identifier of each *cis*-regulatory sequence is indicated. Green: active sequence; red: inactive one. **B)** Similar analysis and representation for the *Ciona robusta Fgf9/16/20* gene, except that the Tcf7 ChIP-seq signal (Oda-Ishii et al., 2016) is shown in yellow. The picture showing the activity of the regulatory sequence at the 16-cell stage is from (Oda-Ishii et al., 2016). **C)** Analysis of chromatin accessibility at the 64-cell stage upstream of the *Phallusia mammillata Nodal* gene. The regions tested by electroporation are indicated (green: active by the larval stage; red: inactive up to the larval stage). Note that NodalPm04, which is evolutionary conserved between *P. mammillata* and *P. fumigata* but not in an open chromatin configuration is inactive by electroporation. **D)** Representative activities at the larval and early gastrula stages of the *Nodal*Pm01, *Nodal*Pm02 and *Nodal*Pm03 sequences. The percentage range of embryos showing staining upon electroporation is indicated. Larval *Nodal*Pm03 staining is found in the progeny of cells stained at the early gastrula stage (b6.5 lineage). Larval *Nodal*Pm01 staining is found both in the progeny of cells stained at the early gastrula stage (TLC, anterior endoderm), as well as in other territories (endodermal strand, muscle).

For each of our test genes, we selected 2 to 7 candidate *cis*-regulatory regions ranging from 209bp to 1316bp (average 525bp) and located up to 4kb upstream from the transcription start site of the gene, within the 5’ UTR, in introns, or immediately downstream of the transcription start site (Figure 6). In most cases, selected regions corresponded to a single reference ATAC-seq peak. Candidate distal cis-regulatory sequences were cloned upstream of a minimal promoter (Table S3) driving NLS-LacZ, to test for enhancer activity. Sequences located immediately upstream of the transcription start site, often coinciding with the H3K4me3 mark, were cloned upstream of the NLS-LacZ reporter, without exogenous minimal promoter, to test for combined enhancer/promoter activity. LacZ activity was monitored at three stages of development: early gastrula (EG), late gastrula (LG) and larva.

In total, we tested the *cis*-regulatory activity of 37 *Phallusia* and 12 *Ciona* candidate open regulatory regions, as well as 11 *Ciona* and *Phallusia* sequences, which were not in an open chromatin state at the 64- and 112-cell stages (Table S4). As an example, the analysis of the *cis*-regulatory sequences controlling the *Phallusia Nodal* gene is presented in Figure 5C, D. Out of the 49 candidate regulatory sequences we electroporated, 73% drove LacZ activity by the larval stage or earlier (Figure 6, blue, green and pink squares), including 16 sequences active at or before the gastrula (Figure 6, blue, green and pink squares outlined by a black line). By contrast, only 3 of 11 closed chromatin sequences (Figure 6, squares outlined with a red line) were active by the larval stage and none of these were active by the gastrula stage. The percentage of open-chromatin regions with *cis*-regulatory activity is thus much larger than the one obtained in previous *cis*-regulatory sequence screens, in which larger (mean 1.7 kb) sequences were chosen at random, irrespective of their chromatin accessibility satus (Harafuji et al., 2002; Keys et al., 2005).

**Figure 6:**
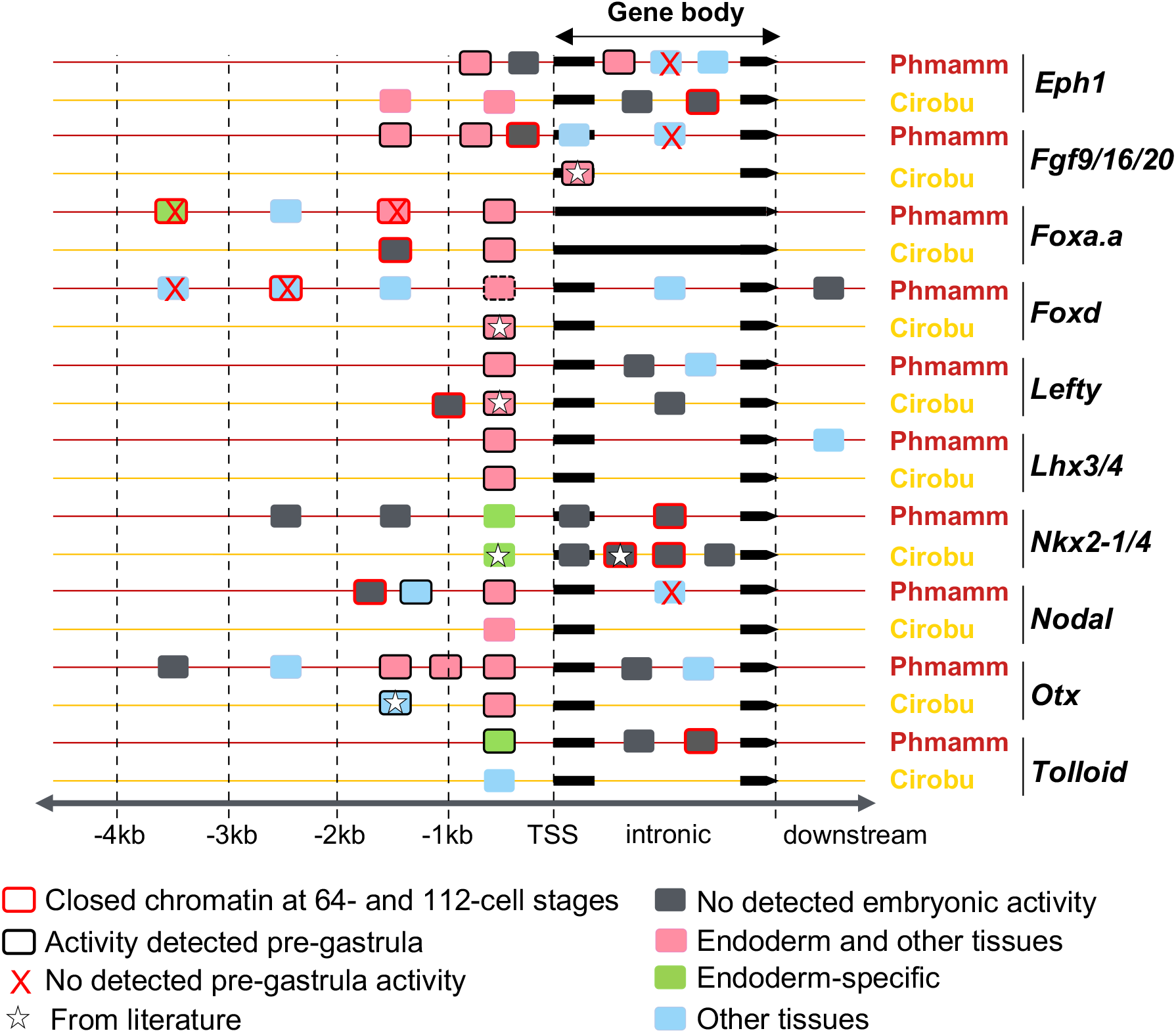
Functional characterisation of the complex regulatory landscapes of early endodermal regulatory genes. Summary of the position and activity of 60 tested sequences and 6 published known regulatory sequences, corresponding to open chromatin regions around orthologous *Phallusia* and *Ciona* genes; the colours represent the spatial tissue specificity of tested regulatory sequences (see Tables S3 and S4 for details). Stars indicate the published known regulatory sequences: *Fgf9/16/20* (ANISEED REG00001080, Oda-Ishii et al., 2016), *Foxd* (ANISEED REG00001115, Oda-Ishii et al., 2016), *Lefty* (ANISEED REG00001118, Oda-Ishii et al., 2016), *Nkx2-1/4* (ANISEED REG00000321 and REG00001175, Fanelli et al., 2003; Doglio et al., 2013) and *Otx* (ANISEED REG00000010, Bertrand et al., 2003). (see Tables S3 and S4 for details). The dotted black line indicates a regulatory sequence with a barely detectable pre-gastrula activity. The schema respects the order of the tested sequences on the chromosome but is not drawn to scale. Only the first and last exons are represented in the gene body.

Interestingly, the timing of onset of activity of an element was related to its position with respect to the transcription start site of the regulated gene. In each of our test genes in both species, an active sequence was located within 500 bp of the transcription start site, identified either by H3K4me3 promoter mark or RNA-seq signal. In at least 15 out of 19 cases, these proximal sequences were active by the early gastrula stage (outlined with a black line on Figure 6). By contrast, only one intronic and 5 distal enhancers, out of a total of 21 active intronic, distal or downstream elements tested at the early gastrula stage, had an early onset of activity (Figure 6).

We conclude that, as expected from global analysis, open chromatin regions detected by ATAC-seq in ascidians are strongly enriched for embryonic *cis*-regulatory elements, confirming that the resource we generated should be of general interest to the community. We also found that ascidian elements with early onset of activity are biased towards a proximal position with respect to the transcription start site and behave as promoter/enhancer elements with spatially restricted activity. It will be interesting to survey whether similar positional bias is also observed in the early embryonic enhancers of other animal species.

### Overlapping temporal and spatial enhancer activity contributes to regulatory landscape complexity

Considering the small number of embryonic territories in ascidian embryos, the *cis*-regulatory landscape in the vicinity of regulatory genes, marked by the presence of COREs, is surprisingly complex. Recent studies in *Drosophila* and vertebrates have revealed that regulatory gene expression is frequently controlled by multiple “shadow” *cis*-regulatory elements with overlapping spatial activities (Barolo, 2012; Cannavò et al., 2016). This partial redundancy was shown to confer robustness of transcriptional regulation to genetic and environmental variations (Frankel et al., 2010; Perry et al., 2010). The *Ciona* genus includes some of the most polymorphic metazoan species known (Leffler et al., 2012) and *Phallusia* and *Ciona* both live in shallow coastal environments subject to strong variations in temperature and salinity. Spatially overlapping enhancer activity may thus be beneficial to ascidians and underlie the unexpected complexity of their early *cis*-regulatory landscapes.

Although a few “shadow” enhancers have been previously identified in *Ciona* (Farley et al., 2016; Fujiwara and Cañestro, 2018; Matsumoto et al., 2008, 2007), the prevalence of enhancers with overlapping activity is unknown. We thus used our collection of active *cis*-regulatory sequences to investigate the presence of regulatory elements with overlapping patterns of activity in the vicinity of our test gene set.

Of the ten genes, five (*Foxa.a, Foxd, FGF9/16/20, Otx* and *Eph1*) were controlled during *Phallusia mammillata* embryogenesis by 2 or 3 *cis*-regulatory sequences with partially overlapping spatial activity (Figure 7), a proportion in keeping with the *Drosophila* situation (Cannavò et al., 2016). By the larval stage (Figure 7A, B), three *Foxa.a cis*-regulatory sequences (*Foxa.a*Pm01, *Foxa.a*Pm03 and *Foxa.a*Pm05) drove activity in the head endoderm and endodermal strand, three *Foxd* sequences (*Foxd*Pm03, *Foxd*Pm05 and *Foxd*Pm06) drove expression in the tail muscles and two *Fgf9/16/20* (*Fgf*Pm03, *Fgf*Pm05) sequences were active in tail muscle cells, B-line notochord and mesenchyme. Analysis of the 12 *Phallusia* cis-regulatory sequences with a pre-gastrula onset of activity identified three additional pairs of regulatory sequences with overlapping spatial and temporal activity (Figure 7C). *Fgf*Pm04 and *Fgf*Pm05 drive early expression in nerve cord precursors. *Otx*Pm01 and *Otx*Pm05 drive early expression in tail muscle and b-line neural tissue. Finally, *Eph1*Pm02 and *Eph1*Pm03 are both active in early endoderm precursors. Based on these examples, ascidian shadow enhancers can be separated by another enhancer (*Otx*Pm01 and *Otx*Pm05) or even by a coding region (*Eph1*Pm02 and *Eph1*Pm03; *Foxd*Pm06 and *Foxd*Pm05).

**Figure 7:**
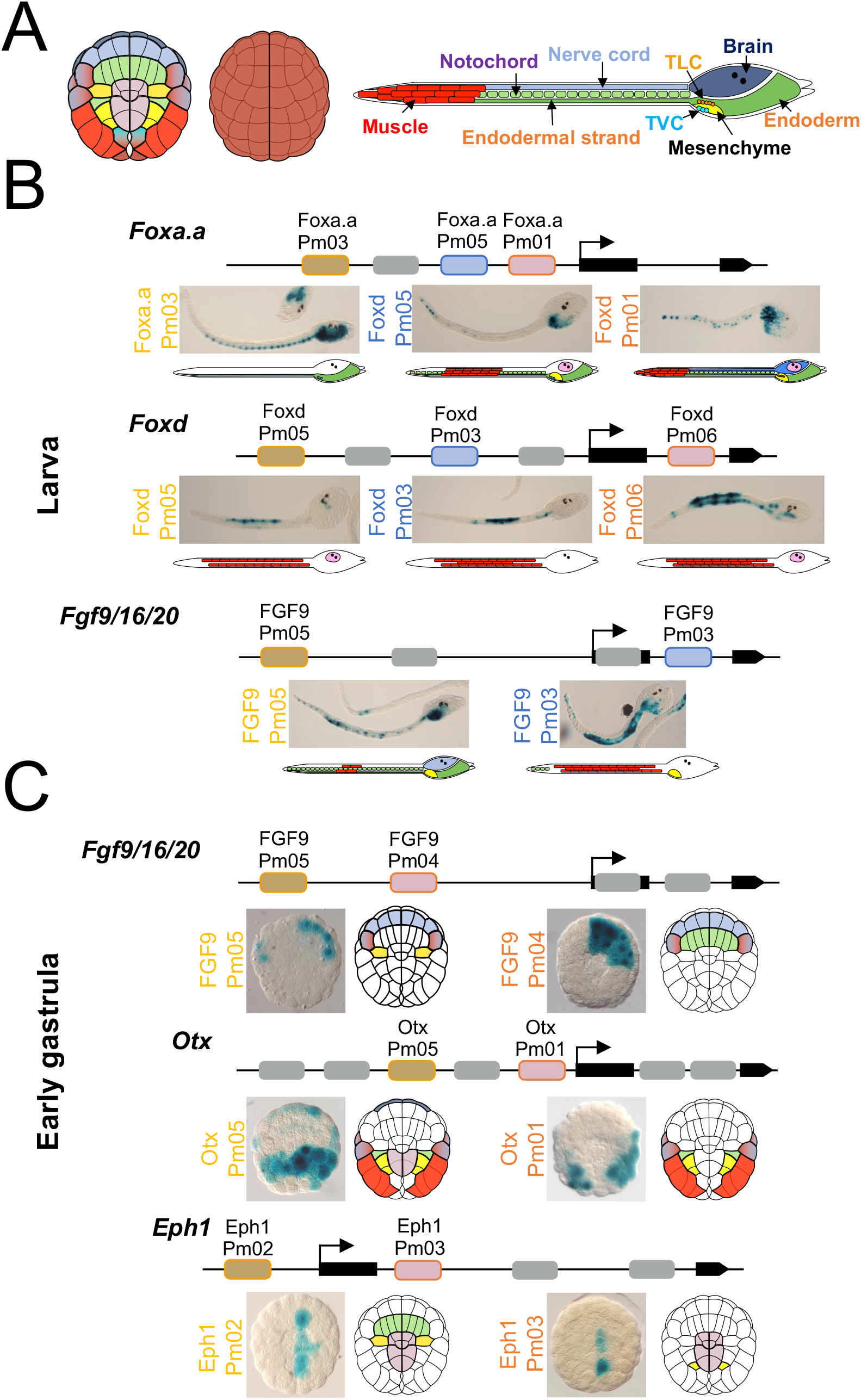
Overlapping temporal and spatial enhancer activity contributes to regulatory landscape complexity. **A)** A schematic of tissues in the *Phallusia mammillata* embryos at the 112-cell stage (left, vegetal view; right, animal view) and larva (epidermal cells were removed to see inner tissues). **B and C)** Schematic of the active regulatory sequences located in the vicinity of the indicated *Phallusia* genes, with their activity at the larval and early gastrula stages, respectively. The coloured rounded rectangles are putative shadow enhancers at the stage considered and the grey rounded rectangles are other tested sequences see Figure 6. The colour of the regulatory sequence indicates to which image it corresponds. Next to the images of the electroporation results are schematics summarising the overall main activity found across the embryos for each construct.

These results suggest that ascidian embryos make frequent use of partially overlapping regulatory sequences, or shadow enhancers, to control their regulatory genes, which may contribute to the complexity of the open chromatin landscape we identified.

### Gastrula *trans*-regulatory activity is temporally and spatially conserved across species

Having extensively characterized the enhancers driving the expression of our ten test genes, we next addressed the evolutionary conservation of their activity between the two species. The evolutionary conservation of gene expression patterns can result from a global conservation of gene regulatory network (GRN) architecture, that is of the combination of transcription factors binding to each cis-regulatory sequence operating within the network. This scenario predicts conservation of the activity of the *cis*-regulatory sequences despite extensive sequence divergence, as observed in insects (Hare et al., 2008), and in a few *Ciona* and *Phallusia* enhancers (Roure et al., 2014). Alternatively, GRNs can be extensively rewired without major impact on regulatory gene expression (Ciliberti et al., 2007; Tolkin and Christiaen, 2016). In this scenario, distinct combinations of transcription factors bind to orthologous *cis*-regulatory sequences causing them to functionally diverge beyond cross-species intelligibility, a situation observed between *Molgula* and *Ciona* species (Lowe and Stolfi, 2018; Stolfi et al., 2014) or between distantly-related echinoderms (Hinman and Davidson, 2007).

To determine which of these two scenarios accounts for the conservation of the early expression of our ten test genes, we individually electroporated 14 *Phallusia* and 3 *Ciona* early *cis*-regulatory sequences active by the gastrula stage into the eggs of the other species and compared the activity of these sequences in the two species at the gastrula stage. 12 of these 17 sequences drove the same patterns of LacZ activity in both species, with an additional 2 elements driving very similar activity (Figure 8A, B) in spite of having diverged beyond alignability between *Phallusia* and *Ciona*. Although we cannot formally exclude that conserved activity results from the binding of different transcription factors in the two species, a parsimonious explanation is that the expression patterns and biochemical activity of the TFs driving the activity of these sequences are conserved between *Ciona* and *Phallusia*. Substantial changes in the activity patterns of the remaining 3 sequences (Figure 8B, *Eph1*Pm02, *Lhx3/4*Ci01, and *Otx*Pm05) could reflect divergence in the specific *trans*-regulatory code driving the activity of these elements between the two species. We note, however, that 11/12 tissue differences in activity across the 17 regulatory sequences are due to the detection of additional activity in *Ciona* (Figure 8B). As electroporation in *Ciona* is more efficient than in *Phallusia* (not shown), these differences could in part reflect an experimental bias, rather than a true evolutionary divergence.

**Figure 8:**
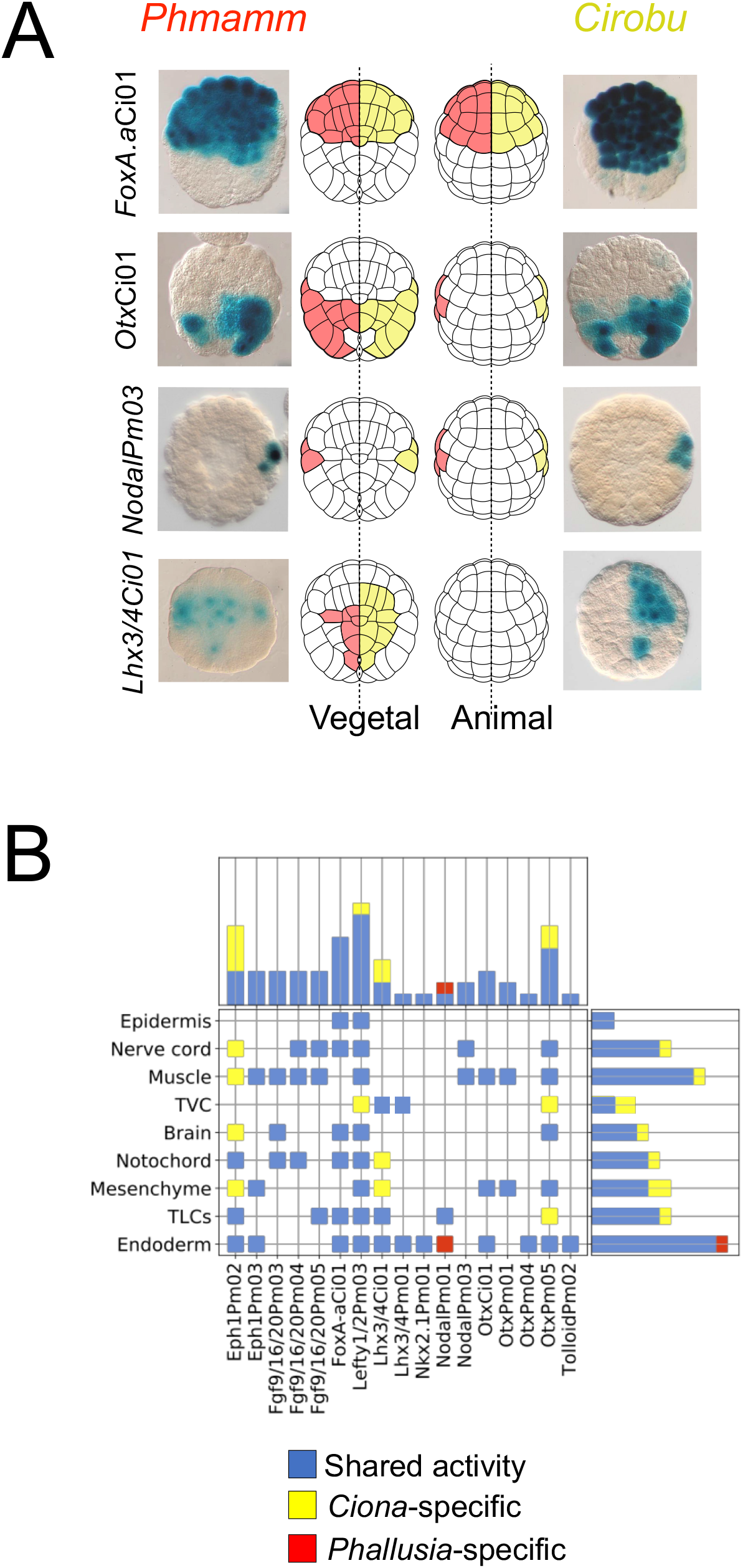
Gastrula *trans*-regulatory activity is temporally and spatially conserved cross-species. **A)** Early gastrula LacZ activity (except *Lhx3/4*Ci01: 64-cell stage) driven in *Phallusia mammillata* and *Ciona intestinalis* by three *Ciona* and one *Phallusia* sequences with conserved activity across the two species. The central schema (left vegetal view, right animal view) summarise on the left embryo half the *Phallusia* activity (pink) and on the right embryo half the *Ciona* activity (yellow). **B)** Summary table of the activity in *Phallusia* and *Ciona* main tissue precursors of 17 sequences as detected by LacZ staining in early or late gastrulae. Blue, red and yellow indicate conserved activity, activity only detected in *Phallusia* and activity only in *Ciona*, respectively. The top box presents a bar graph of the fraction of conserved vs divergent activities of each sequence across all tissues. The side box is a bar graph of the frequency of conserved or divergent activity in each tissue across all sequences.

We conclude that while ascidian *trans*-regulatory landscapes may diverge significantly between the more divergent *Molgula* and *Ciona* ascidian species (Lowe and Stolfi, 2018; Stolfi et al., 2014), early vegetal *trans*-regulatory landscapes appear to be well conserved between the *Phallusia* and *Ciona* species we compared here.

### Collinearity of *Phallusia* and *Ciona cis*-regulatory landscapes around the *FGF9/16/20* and *Otx* genes

The previous section assayed whether the *Ciona* and *Phallusia trans*-regulatory codes are sufficiently close to interpret similarly a given enhancer. In this final section, we conversely addressed whether, despite their sequence divergence, orthologous *Ciona* and *Phallusia* enhancers drive similar activities in a given species, that is the extent of *cis*-regulatory conservation.

For this, we first needed to identify orthologous *Phallusia* and *Ciona* enhancers, a task made difficult by the non-alignability of intergenic and intronic sequences between these two genera. Previous work in insects suggested that the relative position of enhancers for regulatory genes may be under evolutionary constraint, and may thus guide the identification of orthologous enhancers (Cande et al., 2009).

To address whether the global organization of *cis*-regulatory sequences between the two species was conserved, we analysed the genomic architecture of the 9 genes with 1-to1 orthology relationships between the two species. Global gene structure and intron length of orthologs was conserved but local syntenic relationships were frequently broken. Only two genes have conserved both upstream and downstream neighbour genes, *Fgf9/16/20* and *Tolloid*. Two more have retained either their upstream (Otx) or downstream (Foxa.a) neighbour. The remaining five genes (*Nkx2-1/4, Nodal, Lhx3/4, Lefty* and *Eph1*) have non-orthologous upstream and downstream neighbours in the two species.

Although the number of regulatory peaks around *Ciona* and *Phallusia* orthologs was remarkably well conserved (Figure 9A), the position of open chromatin regions was for most genes not directly comparable. Strikingly, however, the upstream and intronic open chromatin landscapes were remarkably conserved between *Ciona* and *Phallusia* for *Fgf9/16/20* and *Otx* (Figure 9B), suggesting that orthologous enhancers can be identified despite the lack of sequence conservation between the two genera. To test this idea further, we compared the activity of three *Phallusia* elements to that of known *Ciona* regulatory sequences (Figure 9B). At the early gastrula stage, the activity of the proximal *Otx*Pm01 element was detected in the posterior mesoderm and b6.5 cell pair progeny. That of the distal *Otx*Pm04 element was detected in the endoderm. The other distal element, *Otx*Pm05, was active in most posterior mesendodermal lineages and in the progeny of the b6.5 and a6.5 cell pairs. These activities were consistent with the reported activities of matching ATAC-seq peaks in *Ciona* (Bertrand et al., 2003). *Otx*Ci01, matching the position of OtxPm01, drove activity in posterior mesoderm, posterior endoderm and b6.5 cell pair progeny. The *Ciona* region matching the position of OtxPm04 was necessary for endoderm activity. Finally, the *Ciona* a-element (REG0000010), corresponding to part of the ATAC-seq peak matching *Otx*Pm05, drove activity in the progeny of the a6.5 and b6.5. Thus, open chromatin regions located in matching positions in the *Ciona* and *Phallusia* intergenic sequences of *Otx* appear to identify orthologous *cis*-regulatory elements.

**Figure 9:**
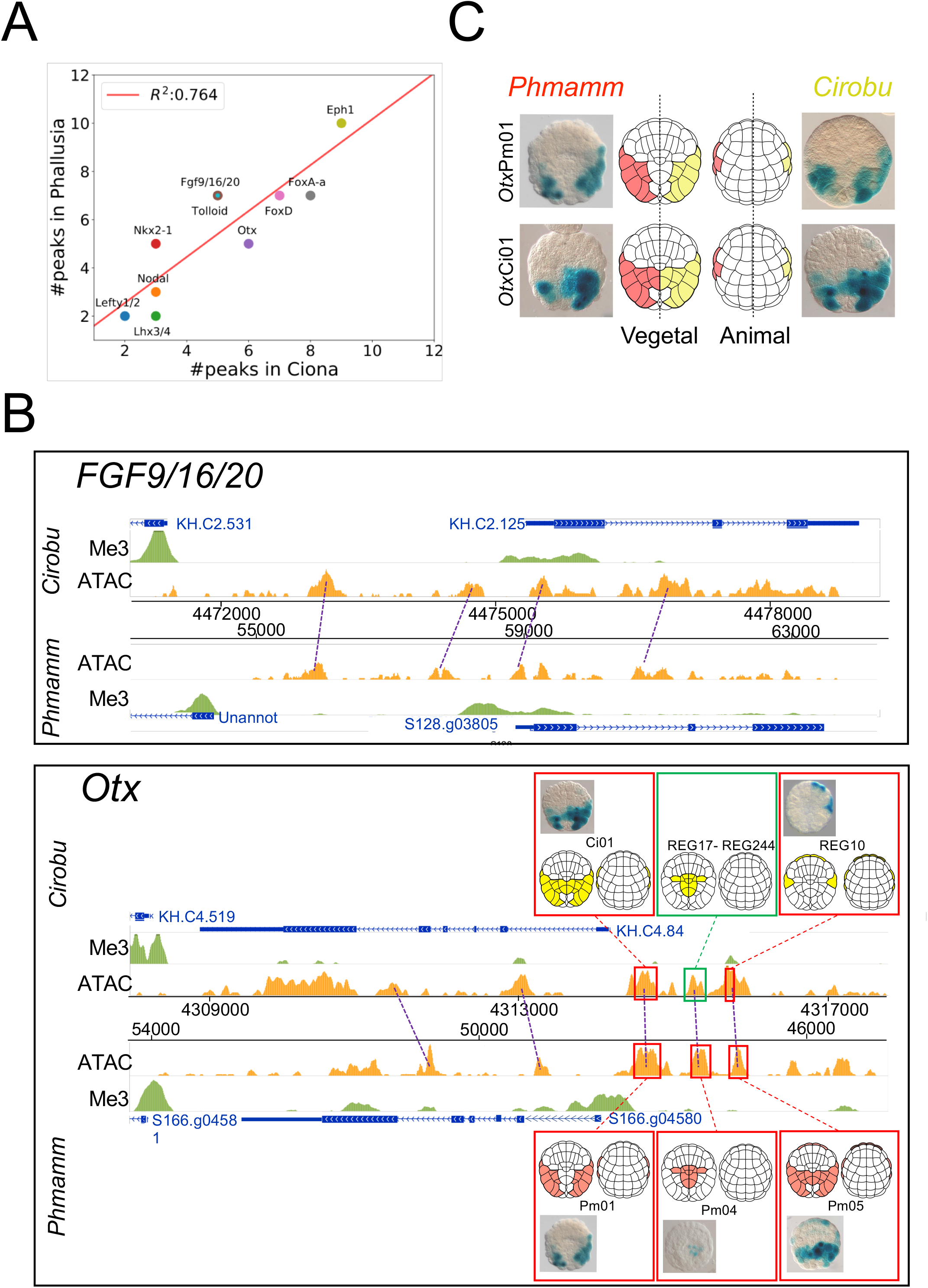
Collinearity of *Phallusia* and *Ciona Otx* and *Fgf9/16/20 cis*-regulatory elements driving qualitatively similar activity. **A)** Graph of the number of open chromatin regions (called ATAC-seq peaks) neighbouring the 10 pairs of orthologous genes studied here. The *x* axis is the number of peaks around the *Phallusia* gene and the *y* axis is the number of peaks around the *Ciona* gene. The regression line is shown in red. The R^2^ value is the coefficient of determination. **B)** Comparison of the open chromatin landscapes at the 64-cell stage of the *Ciona* and *Phallusia Otx* (KH.C4 and MTP2014.S166) and *Fgf9/16/20* (KH.C2 and MTP2014.S128) loci. In the case of *Otx*, the activity of the corresponding peaks tested by electroporation at the early gastrula stage is indicated by a representative picture (which may show mosaic expression) and a schematic representation of the complete pattern of activity. The activity shown represents either the activity of the sequence itself (boxed in red) or an inference of its activity obtained by comparing two constructs differing by this sequence (boxed in green with the identity of the two ANISEED *cis*-regulatory sequences compared). **C)** Early gastrula stage LacZ activity driven in *Phallusia mammillata* and *Ciona robusta* by two orthologous *Otx cis*-regulatory elements. The central schema follows the same logic as Figure 8A.

This orthology did not preclude the evolution of the activity of the enhancers. For example, *Otx*Ci01 drives posterior endodermal expression in both *Ciona* and *Phallusia*, while *Otx*Pm01 does not show this activity in either species (Figure 9C). Thus, orthologous *Fgf9/16/20* and *Otx* early enhancers have retained their relative order and approximate distance to the transcription start site, yet some at least have undergone *cis*-regulatory changes, which partially affected their spatial patterns of activity.

## CONCLUSIONS

In this study, we addressed how *Ciona* and *Phallusia* species, despite their extremely divergent genomes can undergo almost identical early developments. We established ATAC-seq in both species and showed that open chromatin landscapes obey similar rules as in other metazoa (Figures 1, 2): Non-coding open chromatin regions are enriched in GC, highly conserved within each genus, cluster around developmental regulatory genes, mostly transcriptional regulators, and are highly enriched in *cis*-regulatory sequences. Our analysis further suggested that the spatio-temporal dynamics of chromatin accessibility in both species correlates with transcription control of the immediate flanking gene with closest 5’ end (Figure 3). This publicly-accessible ATAC-seq dataset will be of general interest to the ascidian community.

To go beyond a global analysis of regulatory landscapes, we selected ten early *Ciona robusta* regulatory genes with early endodermal expression. We identified the *Phallusia* orthologs and showed that their expression profiles up to the gastrula stages were highly consistent with the *Ciona* data (Figure 4). We identified 39 *cis*-regulatory sequences for these genes, including 17 with activity by the gastrula stage (Figures 5, 6).

Thanks to this dataset, we could reveal features of ascidian transcriptional regulation and its evolution. First, in half of the genes we studied, we found 2 or more enhancers with overlapping spatial activity (Figure 7). This suggests that ascidians make frequent use of shadow enhancers with partially overlapping activity. Given the proposed function of shadow enhancers in ensuring developmental robustness to environmental and genetic stress (Frankel et al., 2010; Perry et al., 2010), it is not surprising that ascidians, which cope with intense environmental variations and a high level of polymorphism, would use this mechanism. The pervasive use of shadow enhancers in the control of regulatory genes may also contribute to the clustering of open chromatin regions in the vicinity of these genes.

Second, both the cross-genera conservation of the expression patterns of the transcriptional regulators we studied and the similar activity of individual early *cis*-regulatory elements in cross-species electroporation assays (Figure 8) argue for a conservation of *trans*-regulatory information between *Ciona* and *Phallusia*, that is a conservation of the combination of transcription factors active in individual embryonic cells. Our results imply that the ascidian *cis*-regulatory code is sufficiently flexible to allow extensive, phenotypically neutral, non-coding sequence divergence within *Phlebobranchia*, thereby limiting the need for compensatory *trans*-regulatory mutations causing Developmental System Drift (True and Haag, 2001). This contrasts with the divergence of *trans*-regulatory information between *Ciona* and the more divergent stolidobranchia *Molgula* (Lowe and Stolfi, 2018; Stolfi et al., 2014).

Finally, we presented evidence that *cis*-regulatory landscapes can be conserved despite extensive sequence divergence (Figure 9). We described that the overall organisation of open chromatin regions at the *Otx* and *Fgf9/16/20* loci is conserved between genera, despite the loss of non-coding sequence alignability. Analysis of the *cis*-regulatory activity of individual open-chromatin regions from both species around Otx, revealed that this conserved organization reflects the presence of orthologous enhancers whose activity, though overall conserved, can slightly diverge. The *cis*-regulatory redundancy ensured by the presence of enhancers with overlapping activity may facilitate this divergence, as the case of divergence we observed involved a posterior endoderm activity redundantly found in two other enhancers at the same locus. The availability of our embryonic ATAC-seq dataset will allow testing on larger datasets whether the evolutionary shuffling of individual redundant regulatory activities between enhancers at a given locus is a general feature of ascidians transcriptional regulation.

## ACKNOWLEDGEMENTS

We thank Dr. A Spagnuolo (Stazione Zoologica Anton Dohrn, Naples, Italy) for the kind gift of *Ciona robusta* genomic DNA, Marc Plays and Philippe Richard for excellent animal husbandry, members of our groups for many discussions and suggestions. We also thank an anonymous referee for her/his encouragements to extend and more thoroughly analyse our initial ATAC-seq dataset during the revision of the manuscript. PL’s lab was supported by grants from the Agence Nationale pour la Recherche (TED project, ANR-13-BSV2-0011). JLG-S’s lab was funded by the European Research Council (ERC) under the European Union’s Horizon 2020 research and innovation program (grant agreement No’s ERC-AdG-LS8), the Spanish Ministerio de Economía y Competitividad (BFU2016-74961-P and BFU2014-55738), the ‘Unidad de Excelencia María de Maetzu 2017-2021’(MDM-2016-0687) to the Department of Gene Regulation and Morphogenesis of CABD), and the Andalusian Government (BIO-396). AM was a PhD student of the University of Montpellier, supported successively by the EpiGenMed Laboratory of excellence and FRM (FDT20160435698). MSM was a PhD student from the University Pablo de Olavide supported by the Spanish Ministerio de Economía y Competitividad, and by an EMBO short-term fellowship to visit PL’s laboratory. DG and SH were supported by TED. UMF and LG were post-doctoral fellows supported by the FRM (SPF20120523969) and HHMI, respectively. PL and CD are members of CNRS and JLG-S is a member of CSIC.

## AUTHORS CONTRIBUTION

Initialisation, funding acquisition and administration of the project: PL and JLG-S. Supervision: PL supervised AM, UMF, SH and CD. AM supervised DG, JLG-S supervised MSM. Investigations: MSM and AM performed the ATAC-seq experiments, AM performed most *in situ* hybridizations, with contributions from UMF. AM, SH and DG performed all cloning and electroporation experiments. Formal analysis: ATAC-seq datasets were computationally processed and analysed by CD and PL. LG participated to the analysis of the electroporation data and to the construction of the figures. AM and PL conceptualized the project and wrote the original draft of the manuscript. All authors reviewed, edited or commented on the manuscript.

## SUPPLEMENTARY TABLE AND FIGURE LEGENDS

**Table S1: Expanded dataset for *Ciona*.**
This table presents the detailed accessibility values for each peak of the reference set for *Ciona* in each condition tested. The data are enriched with RNA-seq statistics for each gene listed. Cells coloured in red correspond to peaks with accessibility scores ≥500 (see methods).

**Table S2: Expanded dataset for *Phallusia*.**
This table presents the detailed accessibility values for each peak of the reference set for *Phallusia* in each condition tested. The data are enriched with RNA-seq statistics for each gene listed. Cells coloured in red correspond to peaks with accessibility scores ≥300 (see methods).

**Table S3: General description of all regulatory sequences described in this article.**
Includes regulated gene, activity class, region Aniseed ID, primers used for amplification, construct information and Aniseed ID.

**Table S4: Spatio temporal regulatory sequence activity following electroporation in *Ciona* and *Phallusia*.**
Red cells indicate the territories of activity.

## REFERENCES

Barolo, S., 2012. Shadow enhancers: Frequently asked questions about distributed cis-regulatory information and enhancer redundancy. BioEssays 34, 135–141. https://doi.org/10.1002/bies.201100121

Bertrand, V., Hudson, C., Caillol, D., Popovici, C., Lemaire, P., 2003. Neural Tissue in Ascidian Embryos Is Induced by FGF9/16/20, Acting via a Combination of Maternal GATA and Ets Transcription Factors. Cell 115, 615–627. https://doi.org/10.1016/S0092-8674(03)00928-0

Brozovic, M., Dantec, C., Dardaillon, J., Dauga, D., Faure, E., Gineste, M., Louis, A., Naville, M., Nitta, K.R., Piette, J., Reeves, W., Scornavacca, C., Simion, P., Vincentelli, R., Bellec, M., Aicha, S.B., Fagotto, M., Guéroult-Bellone, M., Haeussler, M., Jacox, E., Lowe, E.K., Mendez, M., Roberge, A., Stolfi, A., Yokomori, R., Brown, C.T., Cambillau, C., Christiaen, L., Delsuc, F., Douzery, E., Dumollard, R., Kusakabe, T., Nakai, K., Nishida, H., Satou, Y., Swalla, B., Veeman, M., Volff, J.-N., Lemaire, P., 2018. ANISEED 2017: extending the integrated ascidian database to the exploration and evolutionary comparison of genome-scale datasets. Nucleic Acids Res 46, D718–D725. https://doi.org/10.1093/nar/gkx1108

Brozovic, M., Martin, C., Dantec, C., Dauga, D., Mendez, M., Simion, P., Percher, M., Laporte, B., Scornavacca, C., Di Gregorio, A., Fujiwara, S., Gineste, M., Lowe, E.K., Piette, J., Racioppi, C., Ristoratore, F., Sasakura, Y., Takatori, N., Brown, T.C., Delsuc, F., Douzery, E., Gissi, C., McDougall, A., Nishida, H., Sawada, H., Swalla, B.J., Yasuo, H., Lemaire, P., 2016. ANISEED 2015: a digital framework for the comparative developmental biology of ascidians. Nucleic Acids Res 44, D808–D818. https://doi.org/10.1093/nar/gkv966

Buenrostro, J.D., Giresi, P.G., Zaba, L.C., Chang, H.Y., Greenleaf, W.J., 2013. Transposition of native chromatin for fast and sensitive epigenomic profiling of open chromatin, DNA-binding proteins and nucleosome position. Nature Methods 10, 1213–1218. https://doi.org/10.1038/nmeth.2688

Cande, J., Goltsev, Y., Levine, M.S., 2009. Conservation of enhancer location in divergent insects. PNAS 106, 14414–14419. https://doi.org/10.1073/pnas.0905754106

Cannavò, E., Khoueiry, P., Garfield, D.A., Geeleher, P., Zichner, T., Gustafson, E.H., Ciglar, L., Korbel, J.O., Furlong, E.E.M., 2016. Shadow Enhancers Are Pervasive Features of Developmental Regulatory Networks. Current Biology 26, 38–51. https://doi.org/10.1016/j.cub.2015.11.034

Caputi, L., Andreakis, N., Mastrototaro, F., Cirino, P., Vassillo, M., Sordino, P., 2007. Cryptic speciation in a model invertebrate chordate. PNAS 104, 9364–9369. https://doi.org/10.1073/pnas.0610158104

Christiaen, L., Wagner, E., Shi, W., Levine, M., 2009a. Isolation of sea squirt (Ciona) gametes, fertilization, dechorionation, and development. Cold Spring Harb Protoc 2009, pdb.prot5344. https://doi.org/10.1101/pdb.prot5344

Christiaen, L., Wagner, E., Shi, W., Levine, M., 2009b. Whole-mount In situ hybridization on sea squirt (Ciona intestinalis) embryos. Cold Spring Harb Protoc 2009, pdb.prot5348. https://doi.org/10.1101/pdb.prot5348

Ciliberti, S., Martin, O.C., Wagner, A., 2007. Robustness can evolve gradually in complex regulatory gene networks with varying topology. PLoS Comput. Biol 3, e15. https://doi.org/10.1371/journal.pcbi.0030015

Conklin, E., 1905. The organization and cell lineage of the ascidian egg. J. Acad., Nat. Sci. Phila. 13, 1

Cusanovich, D.A., Reddington, J.P., Garfield, D.A., Daza, R.M., Aghamirzaie, D., Marco-Ferreres, R., Pliner, H.A., Christiansen, L., Qiu, X., Steemers, F.J., Trapnell, C., Shendure, J., Furlong, E.E.M., 2018. The cis-regulatory dynamics of embryonic development at single-cell resolution. Nature 555, 538–542. https://doi.org/10.1038/nature25981

Daugherty, A.C., Yeo, R.W., Buenrostro, J.D., Greenleaf, W.J., Kundaje, A., Brunet, A., 2017. Chromatin accessibility dynamics reveal novel functional enhancers in C. elegans. Genome Res. 27, 2096–2107. https://doi.org/10.1101/gr.226233.117

Delsuc, F., Philippe, H., Tsagkogeorga, G., Simion, P., Tilak, M.-K., Turon, X., López-Legentil, S., Piette, J., Lemaire, P., Douzery, E.J.P., 2018. A phylogenomic framework and timescale for comparative studies of tunicates. BMC Biology 16, 39. https://doi.org/10.1186/s12915-018-0499-2

Doglio, L., Goode, D.K., Pelleri, M.C., Pauls, S., Frabetti, F., Shimeld, S.M., Vavouri, T., Elgar, G., 2013. Parallel Evolution of Chordate Cis-Regulatory Code for Development. PLOS Genetics 9, e1003904. https://doi.org/10.1371/journal.pgen.1003904

Duboule, D., 2007. The rise and fall of Hox gene clusters. Development 134, 2549–2560. https://doi.org/10.1242/dev.001065

Fanelli, A., Lania, G., Spagnuolo, A., Di Lauro, R., 2003. Interplay of negative and positive signals controls endoderm-specific expression of the ascidian Cititf1 gene promoter. Developmental Biology 263, 12–23. https://doi.org/10.1016/S0012-1606(03)00397-X

Farley, E.K., Olson, K.M., Zhang, W., Rokhsar, D.S., Levine, M.S., 2016. Syntax compensates for poor binding sites to encode tissue specificity of developmental enhancers. PNAS 113, 6508–6513. https://doi.org/10.1073/pnas.1605085113

Félix, M.-A., Barkoulas, M., 2012. Robustness and flexibility in nematode vulva development. Trends in Genetics 28, 185–195. https://doi.org/10.1016/j.tig.2012.01.002

Fiuza, U.-M., Negishi, T., Rouan, A., Yasuo, H., Lemaire, P., 2018. Nodal and Eph signalling relay drives the transition between apical constriction and apico-basal shortening during ascidian endoderm invagination. bioRxiv 418988. https://doi.org/10.1101/418988

Frankel, N., Davis, G.K., Vargas, D., Wang, S., Payre, F., Stern, D.L., 2010. Phenotypic robustness conferred by apparently redundant transcriptional enhancers. Nature 466, 490–493. https://doi.org/10.1038/nature09158

Fujiwara, S., Cañestro, C., 2018. Reporter Analyses Reveal Redundant Enhancers that Confer Robustness on <Emphasis Type=“Italic”>Cis</Emphasis>-Regulatory Mechanisms, in: Transgenic Ascidians, Advances in Experimental Medicine and Biology. Springer, Singapore, pp. 69–79. https://doi.org/10.1007/978-981-10-7545-2_7

Gaulton, K.J., Nammo, T., Pasquali, L., Simon, J.M., Giresi, P.G., Fogarty, M.P., Panhuis, T.M., Mieczkowski, P., Secchi, A., Bosco, D., Berney, T., Montanya, E., Mohlke, K.L., Lieb, J.D., Ferrer, J., 2010. A map of open chromatin in human pancreatic islets. Nature Genetics 42, 255–259. https://doi.org/10.1038/ng.530

Guignard, L., Fiuza, U.-M., Leggio, B., Faure, E., Laussu, J., Hufnagel, L., Malandain, G., Godin, C., Lemaire, P., 2018. Contact-dependent cell communications drive morphological invariance during ascidian embryogenesis. bioRxiv 238741. https://doi.org/10.1101/238741

Hagey, D.W., Zaouter, C., Combeau, G., Lendahl, M.A., Andersson, O., Huss, M., Muhr, J., 2016. Distinct transcription factor complexes act on a permissive chromatin landscape to establish regionalized gene expression in CNS stem cells. Genome Res. 26, 908–917. https://doi.org/10.1101/gr.203513.115

Harafuji, N., Keys, D.N., Levine, M., 2002. Genome-wide identification of tissue-specific enhancers in the Ciona tadpole. PNAS 99, 6802–6805. https://doi.org/10.1073/pnas.052024999

Hare, E.E., Peterson, B.K., Iyer, V.N., Meier, R., Eisen, M.B., 2008. Sepsid even-skipped enhancers are functionally conserved in Drosophila despite lack of sequence conservation. PLoS Genet 4, e1000106. https://doi.org/10.1371/journal.pgen.1000106

Hinman, V.F., Davidson, E.H., 2007. Evolutionary plasticity of developmental gene regulatory network architecture. Proc. Natl. Acad. Sci. U.S.A 104, 19404–19409. https://doi.org/10.1073/pnas.0709994104

Horie, T., Horie, R., Chen, K., Cao, C., Nakagawa, M., Kusakabe, T.G., Satoh, N., Sasakura, Y., Levine, M., 2018. Regulatory cocktail for dopaminergic neurons in a protovertebrate identified by whole-embryo single-cell transcriptomics. Genes Dev. 32, 1297–1302. https://doi.org/10.1101/gad.317669.118

Hudson, C., Sirour, C., Yasuo, H., 2016. Co-expression of Foxa.a, Foxd and Fgf9/16/20 defines a transient mesendoderm regulatory state in ascidian embryos. eLife Sciences 5, e14692. https://doi.org/10.7554/eLife.14692

Imai, K., Takada, N., Satoh, N., Satou, Y., 2000. (beta)-catenin mediates the specification of endoderm cells in ascidian embryos. Development 127, 3009–20.

Imai, K.S., Hino, K., Yagi, K., Satoh, N., Satou, Y., 2004. Gene expression profiles of transcription factors and signaling molecules in the ascidian embryo: towards a comprehensive understanding of gene networks. Development 131, 4047–58. https://doi.org/10.1242/dev.01270

Imai, K.S., Levine, M., Satoh, N., Satou, Y., 2006. Regulatory blueprint for a chordate embryo. Science 312, 1183–1187. https://doi.org/10.1126/science.1123404

Keys, D.N., Lee, B., Di Gregorio, A., Harafuji, N., Detter, J.C., Wang, M., Kahsai, O., Ahn, S., Zhang, C., Doyle, S.A., Satoh, N., Satou, Y., Saiga, H., Christian, A.T., Rokhsar, D.S., Hawkins, T.L., Levine, M., Richardson, P.M., 2005. A saturation screen for cis-acting regulatory DNA in the Hox genes of Ciona intestinalis. Proc Natl Acad Sci U S A 102, 679–83. https://doi.org/10.1073/pnas.0408952102

Lamy, C., Rothbächer, U., Caillol, D., Lemaire, P., 2006. Ci-FoxA-a is the earliest zygotic determinant of the ascidian anterior ectoderm and directly activates Ci-sFRP1/5. Development 133, 2835–44. https://doi.org/10.1242/dev.02448

Langmead, B., Salzberg, S.L., 2012. Fast gapped-read alignment with Bowtie 2. Nature Methods 9, 357–359. https://doi.org/10.1038/nmeth.1923

Leffler, E.M., Bullaughey, K., Matute, D.R., Meyer, W.K., Ségurel, L., Venkat, A., Andolfatto, P., Przeworski, M., 2012. Revisiting an Old Riddle: What Determines Genetic Diversity Levels within Species? PLOS Biology 10, e1001388. https://doi.org/10.1371/journal.pbio.1001388

Lemaire, P., 2011. Evolutionary crossroads in developmental biology: the tunicates. Development 138, 2143–2152. https://doi.org/10.1242/dev.048975

Li, L., Zhu, Q., He, X., Sinha, S., Halfon, M.S., 2007. Large-scale analysis of transcriptional cis-regulatory modules reveals both common features and distinct subclasses. Genome Biology 8, R101. https://doi.org/10.1186/gb-2007-8-6-r101

Liu, G., Wang, W., Hu, S., Wang, X., Zhang, Y., 2018. Inherited DNA methylation primes the establishment of accessible chromatin during genome activation. Genome Res. https://doi.org/10.1101/gr.228833.117

Lowe, E.K., Stolfi, A., 2018. Developmental system drift in motor ganglion patterning between distantly related tunicates. bioRxiv 320804. https://doi.org/10.1101/320804

Maere, S., Heymans, K., Kuiper, M., 2005. BiNGO: a Cytoscape plugin to assess overrepresentation of gene ontology categories in biological networks. Bioinformatics 21, 3448–3449. https://doi.org/10.1093/bioinformatics/bti551

Martin, M., 2011. Cutadapt removes adapter sequences from high-throughput sequencing reads. EMBnet.journal 17, 10–12. https://doi.org/10.14806/ej.17.1.200

Matsumoto, J., Katsuyama, Y., Ohtsuka, Y., Lemaire, P., Okamura, Y., 2008. Functional analysis of synaptotagmin gene regulatory regions in two distantly related ascidian species. Development, Growth & Differentiation 50, 543–552. https://doi.org/10.1111/j.1440-169X.2008.01049.x

Matsumoto, J., Kumano, G., Nishida, H., 2007. Direct activation by Ets and Zic is required for initial expression of the Brachyury gene in the ascidian notochord. Developmental Biology 306, 870–882. https://doi.org/10.1016/j.ydbio.2007.03.034

Nishida, H., 1987. Cell lineage analysis in ascidian embryos by intracellular injection of a tracer enzyme. III. Up to the tissue restricted stage. Dev Biol 121, 526–41.

Oda-Ishii, I., Bertrand, V., Matsuo, I., Lemaire, P., Saiga, H., 2005. Making very similar embryos with divergent genomes: conservation of regulatory mechanisms of Otx between the ascidians Halocynthia roretzi and Ciona intestinalis. Development 132, 1663–74. https://doi.org/10.1242/dev.01707

Oda-Ishii, I., Kubo, A., Kari, W., Suzuki, N., Rothbächer, U., Satou, Y., 2016. A Maternal System Initiating the Zygotic Developmental Program through Combinatorial Repression in the Ascidian Embryo. PLOS Genetics 12, e1006045. https://doi.org/10.1371/journal.pgen.1006045

Perry, M.W., Boettiger, A.N., Bothma, J.P., Levine, M., 2010. Shadow Enhancers Foster Robustness of Drosophila Gastrulation. Current Biology 20, 1562–1567. https://doi.org/10.1016/j.cub.2010.07.043

Pott, S., Lieb, J.D., 2015. What are super-enhancers? Nature Genetics 47, 8–12. https://doi.org/10.1038/ng.3167

Preissl, S., Fang, R., Huang, H., Zhao, Y., Raviram, R., Gorkin, D.U., Zhang, Y., Sos, B.C., Afzal, V., Dickel, D.E., Kuan, S., Visel, A., Pennacchio, L.A., Zhang, K., Ren, B., 2018. Single-nucleus analysis of accessible chromatin in developing mouse forebrain reveals cell-type-specific transcriptional regulation. Nature Neuroscience 21, 432–439. https://doi.org/10.1038/s41593-018-0079-3

Ristoratore, F., Spagnuolo, A., Aniello, F., Branno, M., Fabbrini, F., Lauro, R.D., 1999. Expression and functional analysis of Cititf1, an ascidian NK-2 class gene, suggest its role in endoderm development. Development 126, 5149–5159.

Roure, A., Lemaire, P., Darras, S., 2014. An Otx/Nodal Regulatory Signature for Posterior Neural Development in Ascidians. PLOS Genetics 10, e1004548. https://doi.org/10.1371/journal.pgen.1004548

Sardet, C., McDougall, A., Yasuo, H., Chenevert, J., Pruliere, G., Dumollard, R., Hudson, C., Hebras, C., Le Nguyen, N., Paix, A., 2011. Embryological methods in ascidians: the Villefranche-sur-Mer protocols. Methods Mol. Biol. 770, 365–400. https://doi.org/10.1007/978-1-61779-210-6_14

Satou, Y., Imai, K.S., Satoh, N., 2001. Early embryonic expression of a LIM-homeobox gene Cs-lhx3 is downstream of beta-catenin and responsible for the endoderm differentiation in Ciona savignyi embryos. Development 128, 3559–70.

Satou, Y., Mineta, K., Ogasawara, M., Sasakura, Y., Shoguchi, E., Ueno, K., Yamada, L., Matsumoto, J., Wasserscheid, J., Dewar, K., Wiley, G., Macmil, S., Roe, B., Zeller, R., Hastings, K., Lemaire, P., Lindquist, E., Endo, T., Hotta, K., Inaba, K., 2008. Improved genome assembly and evidence-based global gene model set for the chordate Ciona intestinalis: new insight into intron and operon populations. Genome Biol 9, R152. https://doi.org/10.1186/gb-2008-9-10-r152

Sharma, S., Wang, W., Stolfi, A., 2018. Single-cell transcriptome profiling of the Ciona larval brain. bioRxiv 319327. https://doi.org/10.1101/319327

Sommer, R.J., 2012. Evolution of Regulatory Networks: Nematode Vulva Induction as an Example of Developmental Systems Drift, in: Evolutionary Systems Biology, Advances in Experimental Medicine and Biology. Springer, New York, NY, pp. 79–91. https://doi.org/10.1007/978-1-4614-3567-9_4

Stern, D.L., Orgogozo, V., 2008. The Loci of Evolution: How Predictable Is Genetic Evolution? Evolution 62, 2155–2177. https://doi.org/10.1111/j.1558-5646.2008.00450.x

Stolfi, A., Lowe, E.K., Racioppi, C., Ristoratore, F., Brown, C.T., Swalla, B.J., Christiaen, L., 2014. Divergent mechanisms regulate conserved cardiopharyngeal development and gene expression in distantly related ascidians. eLife Sciences 3, e03728. https://doi.org/10.7554/eLife.03728

Takahashi, H., Mitani, Y., Satoh, G., Satoh, N., 1999. Evolutionary alterations of the minimal promoter for notochord-specific Brachyury expression in ascidian embryos. Development 126, 3725–3734.

Thomas, S., Li, X.-Y., Sabo, P.J., Sandstrom, R., Thurman, R.E., Canfield, T.K., Giste, E., Fisher, W., Hammonds, A., Celniker, S.E., Biggin, M.D., Stamatoyannopoulos, J.A., 2011. Dynamic reprogramming of chromatin accessibility during Drosophilaembryo development. Genome Biology 12, R43. https://doi.org/10.1186/gb-2011-12-5-r43

Tolkin, T., Christiaen, L., 2016. Rewiring of an ancestral Tbx1/10-Ebf-Mrf network for pharyngeal muscle specification in distinct embryonic lineages. Development 143, 3852–3862. https://doi.org/10.1242/dev.136267

Treen, N., Heist, T., Wang, W., Levine, M., 2018. Depletion of Maternal Cyclin B3 Contributes to Zygotic Genome Activation in the Ciona Embryo. Current Biology 28, 1150–1156.e4. https://doi.org/10.1016/j.cub.2018.02.046

True, J.R., Haag, E.S., 2001. Developmental system drift and flexibility in evolutionary trajectories. Evol. Dev. 3, 109–119.

Tsagkogeorga, G., Cahais, V., Galtier, N., 2012. The Population Genomics of a Fast Evolver: High Levels of Diversity, Functional Constraint, and Molecular Adaptation in the Tunicate Ciona intestinalis. Genome Biol Evol 4, 852–861. https://doi.org/10.1093/gbe/evs054

Wang, W., Niu, X., Jullian, E., Kelly, R.G., Satija, R., Christiaen, L., 2017. A single cell transcriptional roadmap for cardiopharyngeal fate diversification. bioRxiv 150235. https://doi.org/10.1101/150235

Wu, J., Huang, B., Chen, H., Yin, Q., Liu, Y., Xiang, Y., Zhang, B., Liu, B., Wang, Q., Xia, W., Li, W., Li, Y., Ma, J., Peng, X., Zheng, H., Ming, J., Zhang, W., Zhang, J., Tian, G., Xu, F., Chang, Z., Na, J., Yang, X., Xie, W., 2016. The landscape of accessible chromatin in mammalian preimplantation embryos. Nature 534, 652–657. https://doi.org/10.1038/nature18606

Yáñez-Cuna, J.O., Kvon, E.Z., Stark, A., 2013. Deciphering the transcriptional cis-regulatory code. Trends in Genetics 29, 11–22. https://doi.org/10.1016/j.tig.2012.09.007

Zhang, Y., Liu, T., Meyer, C.A., Eeckhoute, J., Johnson, D.S., Bernstein, B.E., Nusbaum, C., Myers, R.M., Brown, M., Li, W., Liu, X.S., 2008. Model-based Analysis of ChIP-Seq (MACS). Genome Biology 9, R137. https://doi.org/10.1186/gb-2008-9-9-r137

